# Soil chemical and microbial gradients determine accumulation of root exuded secondary metabolites and plant-soil feedbacks in the field

**DOI:** 10.1101/2023.06.09.544436

**Authors:** Valentin Gfeller, Selma Cadot, Jan Waelchli, Sophie Gulliver, Céline Terrettaz, Lisa Thönen, Pierre Mateo, Christelle A.M. Robert, Fabio Mascher, Thomas Steinger, Moritz Bigalke, Matthias Erb, Klaus Schlaeppi

**Affiliations:** Institute of Plant Sciences, University of Bern, 3013 Bern, Switzerland; Department of Environmental Sciences, University of Basel, 4056 Basel, Switzerland; Division of Plant Breeding, Agroscope, 1260 Nyon, Switzerland; Division of Plant Protection, Agroscope, 1260 Nyon, Switzerland; Institute of Geography, University of Bern, 3012 Bern, Switzerland

**Keywords:** Plant-soil feedback, crop rotation, secondary metabolites, plant-microbe interactions, wheat, maize, environmental gradient, soil chemistry

## Abstract

**Introduction:** Harnessing positive plant-soil feedbacks via crop rotations is a promising strategy for sustainable agriculture. Plants can influence soil properties including microbes by exuding specialized metabolites. However, the effects are often context dependent and variable. If and how local soil heterogeneity may explain this variation is unknown. Benzoxazinoids are specialized metabolites that are released in high quantities by cereals such as wheat and maize. Benzoxazinoids can alter rhizosphere microbiota and the performance of plants subsequently growing in the exposed soils and are thus an excellent model to study agriculturally relevant plant-soil feedbacks in the field, and to assess how soil factors affect their outcome.

**Materials & methods:** To understand the importance of local variation in soil properties on benzoxazinoid-mediated plant-soil feedbacks, we conditioned plots with wild-type maize and benzoxazinoid-deficient *bx1* mutant plants in a grid pattern across an arable field. We then grew winter wheat across the entire field in the following season. We determined accumulation of benzoxazinoids, root-associated microbial communities, abiotic soil properties and wheat performance in each plot. We also determined benzoxazinoid conversion dynamics in a labelling experiment under controlled conditions, and then assessed associations between soil chemical variation and benzoxazinoid-mediated plant-soil feedbacks.

**Results:** Across the field, we detected a marked gradient in soil chemical and microbial community composition. This gradient resulted in significant differences in benzoxazinoid accumulation. These differences were explained by differential benzoxazinoid degradation rather than exudation. Benzoxazinoid exudation modulated alpha diversity of root and rhizosphere bacteria and fungi during maize growth, but not during subsequent wheat growth, while the chemical fingerprint of benzoxazinoid accumulation persisted. Averaged across the field, we detected no significant feedback effects of benzoxazinoid conditioning on wheat performance and defence, apart from a transient decrease in biomass during vegetative growth. Closer analysis however, revealed pronounced feedback effects along the chemical and microbial gradient of the field, with effects gradually changing from negative to positive along the gradient.

**Conclusion:** Overall, this study revealed that plant-soil feedbacks differ in strength and direction within a field, and that this variation can be explained by standing chemical and microbial gradients, which strongly affect benzoxazinoid accumulation in the soil. Understanding within-field soil heterogeneity is crucial for the future exploitation of plant-soil feedbacks in sustainable precision agriculture.

## Introduction

Plants influence the soil they grow in, which in turn influences the performance of subsequent plants. In crop production, designing a suitable crop rotation makes use of positive plant-soil feedbacks (van der Putten *et al.*, 2013). Identifying and exploiting the mechanisms of plant-soil feedbacks in crop rotations has been proposed as a promising tool to promote sustainable agriculture by leveraging agroecological effects (Mariotte *et al.*, 2018). A key challenge in this context is the spatial and temporal variability of plant-soil feedbacks in the field (Smith-Ramesh & Reynolds, 2017).

Plant-soil feedbacks are attributed to a number of different mechanisms, including changes in mutualist and pathogen abundance, microbiome composition, nutrient availability, and other soil chemical properties (Bever *et al.*, 2012; Bennett & Klironomos, 2019; Pineda *et al.*, 2020). These drivers can affect germination, plant performance (Tawaha & Turk, 2003; van der Putten *et al.*, 2013), as well as pathogen and herbivore resistance (Kos *et al.*, 2015b; Ma *et al.*, 2017; Pineda *et al.*, 2020). Given that all these factors are highly heterogeneous, one can expect strong context dependency and spatiotemporal variation in the resulting feedback effects.

Plant-associated microbial communities have gained attention as drivers of plant-soil feedbacks in the past (Bever *et al.*, 2012). An important mechanism in how plants shape their microbiome is through root exudates, defined as the secretion of primary and secondary metabolites to the surrounding soil (Pang *et al.*, 2021). Prominent examples of secondary metabolites involved in structuring root or rhizosphere microbiomes are benzoxazinoids, coumarins, flavones, and triterpenes (Hu *et al.*, 2018b; Huang *et al.*, 2019; Stringlis *et al.*, 2019; Voges *et al.*, 2019; Yu *et al.*, 2021). Changes in root-associated microbiomes can in turn increase plant growth and defence (Berendsen *et al.*, 2012; Pieterse *et al.*, 2014).

Spatial and temporal variation in plant-soil feedbacks has been studied mostly for soil nutrients, temperature (Smith-Ramesh & Reynolds, 2017; Long *et al.*, 2019), drought (Fry *et al.*, 2018), and the interaction of abiotic factors with soil biota (Kaisermann *et al.*, 2017; Long *et al.*, 2019). Further, local soil biotic communities, represent key determinants of plant-soil feedbacks across time and space (Revillini *et al.*, 2016; Bennett *et al.*, 2017). How environmental heterogeneity affects plant-soil feedbacks driven by exuded root metabolites is largely unknown, and local (i.e., within-field) variation in the underlying dynamics has not been studied so far. Furthermore, the dynamic interplay between exuded metabolites and their degradation or metabolization by rhizosphere microbial communities can be expected to add to the variation in plant-soil feedbacks.

Benzoxazinoids, a class of secondary metabolites common in grasses including maize and wheat, have been shown to be bioactive in many ways (Niemeyer, 2009). They are long known to be involved in allelopathy and defence against insects and pathogens (Niemeyer, 2009; Schandry & Becker, 2020). More recently they have been shown to alleviate plant growth suppressive effects provoked by preceding plants (Gfeller *et al.*, 2023). They can also chelate iron and aluminium (Hu *et al.*, 2018a; Zhou *et al.*, 2018; Zhao *et al.*, 2019). In the past few years benzoxazinoids have repeatedly been shown to shape root-associated microbiomes (Hu *et al.*, 2018b; Cotton *et al.*, 2019; Kudjordjie *et al.*, 2019; Cadot *et al.*, 2021b). Maize roots predominantly excrete DIMBOA-Glc, HDMBOA-Glc, and DIMBOA and these benzoxazinoid compounds are rapidly converted into MBOA in soil (Hu *et al.*, 2018b). Ultimately, soil microorganisms can further metabolize MBOA (half-live: days to weeks) to AMPO (weeks to months) and thereby alter their availability in soils (Macías *et al.*, 2004; Etzerodt *et al.*, 2008). So far, benzoxazinoid-dependent plant-soil feedbacks have been shown for maize-maize and maize-wheat cropping sequences in the greenhouse (Hu *et al.*, 2018b; Cadot *et al.*, 2021a) and in the field (Gfeller *et al.*, 2022). In the relatively homogenous experimental field, benzoxazinoid exudation by maize resulted in an increase in wheat yield. If and how such feedbacks act under more heterogeneous conditions is largely unknown.

In this study, we investigated in a maize-wheat crop rotation how soil heterogeneity influences benzoxazinoid-mediated plant-soil feedbacks. We set up a two-year field experiment where we first grew maize to condition the soil followed by winter wheat to score the feedbacks (**Fig. 1**). The soil was conditioned either by benzoxazinoid-producing wild-type maize or by benzoxazinoid-deficient *bx1* mutant plants, and we then assessed feedbacks in 20 replicate plots within the field. Feedback effects were then analysed taking the gradient in soil chemistry present in the field into account. Detailed measurements of benzoxazinoid accumulation and changes in soil microbiota were used to determine to what extent these factors interact with soil heterogeneity to explain the observed variation in plant-soil feedbacks. Overall, our results show a high context dependency of secondary metabolite-mediated plant-soil feedbacks. Understanding such context dependencies is crucial to successfully employ the concept of plant-soil feedbacks in crop rotations.

**Figure 1.**
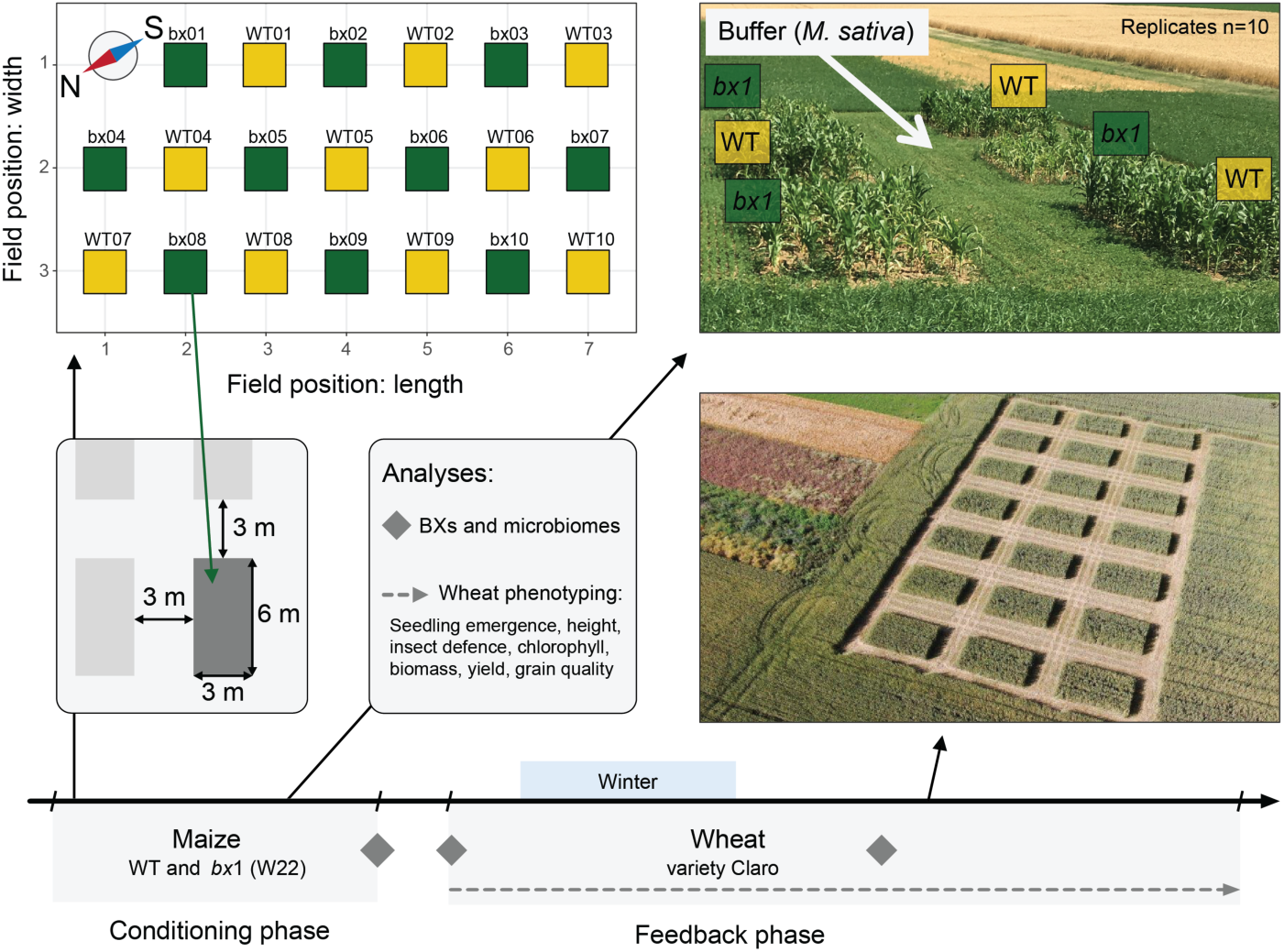
Experimental setup. To examine the effect of maize benzoxazinoid soil conditioning on subsequent wheat growth, defence, yield, and grain quality we conducted a two-year field experiment in Changins, Switzerland. First, wild-type (WT) and benzoxazinoid-deficient *bx1* mutant plants of the maize line W22 were grown on 10 plots each (plot dimensions 3 m x 6 m). As a buffer, *Medicago sativa* was grown between maize plots. After maize harvest, the winter wheat variety Claro was sown. Wheat growth and defence were intensively phenotyped throughout cultivation. At harvest, yield was determined, and grain quality was analysed. Soil benzoxazinoid concentrations and microbiomes were analysed at maize harvest, wheat sowing, and wheat growth. For microbiome analysis roots, rhizospheres, and the soils surrounding the plants were sampled, except for the time point at wheat sowing, where only soil was present. For more details, please refer to the method section.

## Material and Methods

### Plant material

Wild-type maize (*Zea mays*) and the corresponding benzoxazinoid-deficient *bx1* transposon insertion mutant (referred to as *bx1*) of the inbred line W22 (Tzin *et al.*, 2015) were planted in the field for soil conditioning. To subsequently test benzoxazinoid feedbacks on wheat, we grew the winter wheat variety CH Claro (Agroscope/DSP; referred to as Claro), a variety of top baking quality class (Strebel *et al.*, 2022), and recommended cultivar for cultivation in Switzerland.

### Field experiment

The field experiment was conducted in Changins (Nyon, Switzerland) on a field at Agroscope (Parcel 29, 46°23’’58’N, 6°14’’25’E; **Fig. 1**) with neutral pH clay loam soil. It consisted of a maize-wheat rotation, maize being the conditioning crop and wheat the feedback crop. The field consisted of 10 plots with wild-type maize alternating with 10 plots with *bx1* mutants. Each plot was 6 m long and 3 m wide and separated from each other by 3 m of buffer of alfalfa (*Medicago sativa*). Maize was grown from Mai to September 2018 followed by wheat from October to July 2019. Exact timing, field management and previous crops are documented in the **Supplementary Methods**.

### Sample collection

To investigate soil benzoxazinoid content and to test for effects on the microbes, we collected samples at the end of maize growth and at the beginning and during wheat growth. After maize growth we sampled root, rhizosphere, and soil samples. For that, a soil core (20 x 20 x 20 cm) from one randomly selected plant per plot was excavated and used for chemical and microbial analysis (n = 10). At the beginning of the feedback phase, we sampled again soil, one day after wheat sowing. At 10 randomly selected positions on each plot, soil cores of the top 20 cm were taken with a 17 mm diameter soil sampler. Samples of each plot were combined and further processed for chemical and microbial analysis (n = 10). We sampled again root, rhizosphere, and soil during wheat growth. From 3 randomly selected plants per plot, the root system (7 x 7 cm wide, 12 cm deep) was excavated and pooled for chemical and microbial analysis (n = 10). To study within-field variation of soil parameters, soil was sampled after the experiment ended. Soil was taken from 5 randomly selected positions at a depth of 5-20 cm, resulting in a total of 2 kg of pooled soil per plot (n =20) for chemical analysis.

### Wheat phenotyping

#### Emergence and vegetative growth

One month after sowing, we counted all emerged seedlings along 1 m of three randomly selected wheat rows per plot to determine possible benzoxazinoid soil conditioning effects on wheat emergence. The sum of all counted seedlings per plot was taken and wheat emergence per area was calculated. To determine wheat vegetative growth, we measured plant chlorophyll content, height, and biomass accumulation. Chlorophyll content of 15 randomly selected flag leaves per plot was measured with a SPAD-502 chlorophyll meter (Konica Minolta, Japan) and the average value was taken for statistical analysis. Height of 5 randomly selected wheat plants per plot was determined and averaged for statistical analysis. Aboveground biomass accumulation was harvested from 2x 1 m of wheat row per plot at ground level. Dry weight was determined after the plant material was dried at 80 °C until constant weight. The obtained data was used to calculate biomass accumulation per area.

#### Insect infestation

Insect infestation was evaluated by counting the number of *Oulema melanompus* larvae during wheat growth. The number of larvae was scored by randomly selecting 5x 0.5 meter of wheat row on each plot and counting larvae present on the flag leaves. In parallel, the number of tillers of these plants was recorded to calculate the number of larvae per plant.

#### Insect performance

To further evaluate plant defence, we assessed insect performance on detached wheat leaves. For that, we collected 2 randomly selected wheat plants per plot and stored them in a zip-lock plastic bag moistened with a wet cotton pad at 4 °C, to draw upon as required throughout the insect performance assay. Two transparent solo cups (4 cm height and 3.5 cm diameter) per plot were equipped with a wet filter paper, and the top 6 cm of the youngest fully developed leaf was placed inside. *Spodoptera littoralis* larvae were reared on an artificial diet until used in the bioassay. One healthy 3^rd^ instar larva was pre-weighed on a microbalance and placed on the wheat leaf before closing the solo cup with an air permeable lid. Leaves were moistened daily and renewed on day 2 and 4 of the assay to assure excess food for all larvae. Larval performance was evaluated by weighing them after one week of feeding. Larvae weight gain per day was calculated (weight end - weight start / number of days feeding * 100) and the mean of the two replicates per plots was used for statistical analysis.

#### Harvest

To estimate final plant biomass, we collected again 2x 1 m of a wheat row on each plot 12 days before harvest. The plant material was dried at 80 °C until constant weight and dry biomass determined. The number of tillers was counted and plant density and aboveground biomass per tiller was calculated. The wheat was harvested once the kernels were ripe (14 % humidity). Nine m^2^ per plot were harvested with a compact plot combine harvester (Quantum, Wintersteiger), and kernel weight per plot was determined. A subset of these kernels was taken for analysing agronomic kernel quality and food quality related parameters (described in the **Supplementary Methods**).

### Soil chemistry

To confirm the gradient of soil chemistry in the field, freshly collected soil was sent to LBU Laboratories (Eric Schweizer AG, Thun, Switzerland) and analysed with different extraction methods: water (H_2_O), ammonium acetate EDTA (AAE), and carbon dioxide saturated water (CO_2_). H_2_O extracts are a proxy for plant available nutrients, AAE extracts represent nutrients available through plant chelation mechanisms and CO_2_ extracts are a common extraction procedure for magnesium, phosphorus, and potassium. In addition, total iron was extracted in nitric acid (HNO_3_) and quantified with inductively coupled plasma mass spectrometry (ICP-MS) as previously described (Cadot *et al.*, 2021b).

### Benzoxazinoid analysis

The extraction of benzoxazinoids and their degradation products from soil as well as the analytical protocol are detailed in the **Supplementary Methods**, where we also list the measured compounds with their abbreviations, chemical name and where they were sourced from. Benzoxazinoids and degradation products were analysed with an Acquity UHPLC system coupled to a G2-XS QTOF mass spectrometer (Waters AG, Bade-Dättwil, Switzerland) as previously described (Gfeller *et al.*, 2022).

### Benzoxazinoid exudation experiment

To evaluate if maize benzoxazinoid exudation or degradation depends on soil chemical parameters, we performed a climate chamber experiment comparing soils from the north (N: plots WT07/bx04) and south (S: plots WT10/bx07) ends of the field. The soil was sieved (10 mm mesh size), and used to fill 130 mL pots before a wild-type (W22) maize seed was sown (n = 10). Plants were grown in walk-in climate chambers under controlled conditions (day/night: 14/10h; temperature: 22°C/18°C; light 550 µmol m^-2^s^-1^; humidity: 60%) and fertilized twice a week with 10 mL of nutrient solution (0.4% (w/v); Plantactive Typ K, Hauert) supplemented with iron (1‰ (w/v); Sequestrene rapid, Maag) twice a week. All plants were randomized weekly, watered as needed and harvested after 3 weeks. First, benzoxazinoid exudation was measured by taking a given plant out of the pot, gently removing the soil of the root system at the very bottom from 4 randomly selected root tips and rinsing 2 cm of the root tips 4x with 100 µL of sterile water. Immediately after, 60 µL of this suspension was added to 140 µL of pure acidified MeOH resulting in MeOH/H_2_O (70:30 v/v; 0.1% formic acid). After centrifugation for 10 minutes at 19’000 x g the supernatant was stored at -20 °C prior to analysis of benzoxazinoids (as described above). Second, roots were cut at soil level, cleaned off adhering soil with water and dried at 80 °C until constant weight. Dry biomass was subsequently determined on a microbalance. Third, the remaining soil in the pot was homogenized, passed through a 5 mm test sieve, 25 mL were put in a 50 mL centrifuge tube, and stored at -80 °C. Benzoxazinoid extraction and measurement of this soil was done as described above. For statistical analyses the benzoxazinoid concentrations in the soil were corrected for differences in root dry weight.

### Benzoxazinoid degradation experiment

To evaluate possible differences in the benzoxazinoid degradation in soils at both ends of the field (N: plots WT07/bx04; S: plots WT10/bx07/WT03), we performed a degradation experiment with labelled deuterated DIMBOA-*d_3_* under controlled conditions in the laboratory. A 10 mL (≈10 mg) aliquot of this soil was mixed with 10 mL of sterile water in a 50 mL centrifuge tube and blended with a Polytron (30 s at 15’000 rpm), to obtain a homogenous suspension (n = 4 per soil). The soil acidity was between pH 6.96 and pH 7.13, therefore we used a phosphate buffer at pH 7 for the negative and no soil controls. Six mL of soil suspension or buffer were transferred into 14 mL culture tubes and incubated at 22 °C in a thermoshaker (at 150 rpm) under oxic conditions. We let the soil acclimate for 3 days before the DIMBOA-*d_3_* was added. DIMBOA-*d_3_* was dissolved in autoclaved deionized water and added to each culture tube (except negative controls) to obtain a final concentration of 30 µg/mL (≈140 µmol/L). To elucidate the kinetics of benzoxazinoid degradation, we sampled the reaction mixes after 1 min, 7.5 min, 15 min, 1 hour, 4 hours, 1 day, and 4 days. For each sampling point, 300 µL reaction mix was pipetted into a 1.5 mL centrifuge tube containing 700 mL acidified MeOH to result in MeOH/H_2_O (70:30 v/v; 0.1% formic acid). The suspension was vigorously vortexed and stored at -80 °C. Once all samples were collected the tubes were thawed, soil particles were removed by centrifugation (20 min, 19’000 x g, 4 °C), the supernatant was filtered (Target2TM, Regenerated Cellulose Syringe Filters, pore size: 0.45 µm; Thermo Scientific) and stored in a glass vial at -20 °C until analysis. Benzoxazinoids were analysed as described above.

### Microbiota profiling

The **Supplementary Methods** contains the details of sample preparation, DNA extraction and the PCR protocol for microbiota profiling. In brief, the collected samples were processed as previously described (Gfeller *et al.*, 2022) and DNA was extracted using the Spin Kit for Soil (MP Biomedical, USA), following the instructions of the manufacturer. After DNA quantification using the AccuClear Ultra High Sensitivity dsDNA Quantitation Kit (Biotium, USA), bacterial and fungal libraries were constructed largely following our two-step PCR profiling protocol described earlier (Gfeller *et al.*, 2022). Briefly, bacterial and fungal profiles were based on PCR primer pairs 799-F (Chelius & Triplett, 2001) and 1193-R (Bodenhausen *et al.*, 2013) and ITS1-F (Gardes & Bruns, 1993) and ITS2-R (White *et al.*, 1990), respectively. PCR products were equimolarly pooled followed by ligation of the Illumina adapters by the Next Generation Sequencing Platform at University of Bern, where they were subsequently sequenced on a MiSeq instrument (v3 cell, paired-end 2 × 300 bp; Illumina, USA). The raw sequencing data is available from the European Nucleotide Archive (http://www.ebi.ac.uk/ena) with the study accession PRJEB59165 (sample IDs ERS14468209 and ERS14468210). The sequencing data was processed as previously described (Gfeller *et al.*, 2022). In short, the bioinformatic pipeline includes the following tools: *FastQC* and *cutadapt*, (Andrews, 2010; Martin, 2011), *DADA2* (Callahan *et al.*, 2016) and databases *SILVA* v.132 (Quast *et al.*, 2013; Callahan, 2018) and *UNITE* v8.1 (Nilsson *et al.*, 2019) for bacterial and fungal taxonomy. Code and the meta data are available on GitHub (https://github.com/PMI-Basel/Gfeller_et_al_Changins_field_experiment).

### Statistical analyses

The soil chemical data was analysed and visualised using principal component analysis (PCA, *FactoMineR*; Lê *et al.*, 2008). PC axes were extracted for further analysis. First, the PC axes were used to check for correlations of soil parameters with the field position. Second, in further analysis the first PC axis, referred to as ‘soil chemistry PC1’, was factored in the linear models to account for variation explained by soil parameters.

The microbiota analyses are detailed in the **Supplementary Methods.** In brief, microbiomes were analysed using the v*egan* package (Oksanen *et al.*, 2020) for rarefaction analysis, unconstrained Principal Coordinate Analysis (PCoA, Bray-Curtis) and permutational analysis of variance (PERMANOVA, 999 permutations; Bray-Curtis). The gradient in soil chemistry was included in all models taking PC1 as a cofactor. Further analyses of alpha and beta diversity were performed using the R package *phyloseq* (McMurdie & Holmes, 2013).

Differences in concentrations of soil benzoxazinoids and their degradation products between the two maize genotypes at the end of the conditioning phase, at wheat sowing, and during wheat growth were tested by Wilcoxon rank-sum tests and false discovery rate (FDR) corrected *p* values were reported (Benjamini & Hochberg, 1995), followed by correlation analysis to test for associations between benzoxazinoid concentrations and soil chemistry (using PC1).

Wheat growth and defence related data were analysed by Analysis of Variance (ANOVA). Homoscedasticity and normal distribution of error variance was checked visually. For plant phenotypes two different statistical analyses were applied: (i) overall benzoxazinoid conditioning effects, effects of the chemical gradient, and the interaction between the two variables were tested with a linear model: lm(phenotype ∼ soil conditioning * soil chemistry PC1); (ii) to test for *local* benzoxazinoid-dependent plant-soil feedbacks, we calculated the log-response ratio (LRR) for every plot. This was calculated with the following formulae for wild-type (*log(local value/surrounding mean)*) and *bx1* (*log(surrounding mean/local value*)) conditioned plots, where *log()* is the natural logarithm, the *local value* is the realised value of a certain phenotype on the plot of interest, and the *surrounding mean* is the mean of all adjacent plots of the opposite treatment (**Fig. S1**). The LRRs were then used to test associations between soil parameters and the direction and strength of the feedback, where positive LRRs indicate positive benzoxazinoid plant-soil feedbacks.

In the greenhouse experiment and laboratory degradation experiments, differences between the two soil origins (S, N) were tested by means of Welch’s two-sample t-tests and false discovery rate (FDR) corrected *p* values were reported (Benjamini & Hochberg, 1995).

Analyses were conducted using the open-source software R (R Core Team, 2021). Data management and visualisation was facilitated with the *tidyverse* packages (Wickham *et al.*, 2019). All code for statistical analysis and visualization and the corresponding data can be downloaded from GitHub (https://github.com/PMI-Basel/Gfeller_et_al_Changins_field_experiment).

## Results

### Strong chemical gradient within an experimental field

To assess the soil chemical properties across our experimental field, we measured pH and nutrients in water (H_2_O), carbon dioxide (CO_2_) and ammonium acetate EDTA (AAE) extracts on the 20 experimental plots arranged in a grid pattern (**Fig. 1**). Principle Component Analysis (PCA) revealed a strong chemical gradient. Axis 1, associated mostly with the length of the field, explained 60% of the chemical variation (**Fig. 2A**, **Fig. S2**), while the axis 4 (associated largely with width of the field) explained 7% of the variation. Overall, we observed the chemical gradient running roughly north-south in a diagonal across the field (**Fig. 2B**). This gradient was also apparent when looking at individual soil nutrients (**Fig. S3**). It was characterized by elevated levels of Ca (H_2_O extracts), K (CO_2_ extracts) and Mn, P and Bo (all AAE extracts) towards the northern corner of the field and elevated levels of water-soluble iron and magnesium (all extracts) towards the southern. To account for this chemical gradient in the field, we included the PCA axis 1, referred to as soil chemistry PC1, as covariable in all downstream analyses.

**Figure 2.**
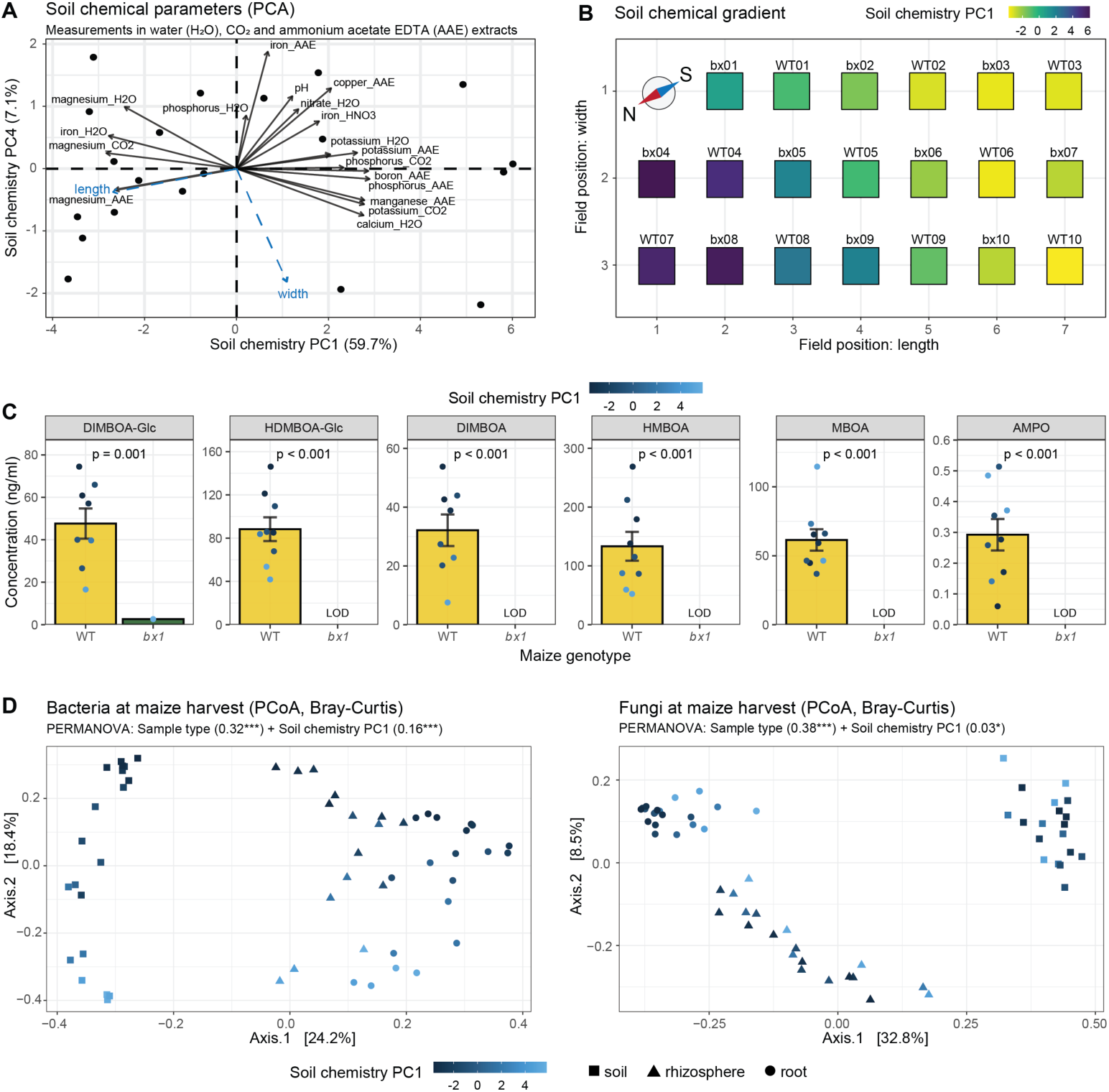
Within-field variation of soil chemistry, benzoxazinoids and microbiota. (A) Principal Component axes 1 (PC1) and 4 (PC4) of soil chemistry PCA are shown. Individual samples (circles), soil parameters (arrows), and direction of field width and length (blue arrows) are included. (B) Field map showing values of soil chemistry PC1 across the field. (C) Soil benzoxazinoid concentrations collected on plots conditioned by wild-type (WT) or benzoxazinoid-deficient *bx1* mutant plants in ng/mL soil (Means ± SE). Statistical significance was calculated by Wilcoxon rank-sum tests and *p* values were corrected for multiple testing. (D) Unconstrained Principal Coordinate Analysis (PCoA) using Bray-Curtis distances of bacterial (left) and fungal (right) communities in soil, rhizosphere, and root samples. *R^2^* and significance level of PERMANOVA on Bray-Curtis distances for bacteria and fungi are shown. Levels of significance: p < 0.001 ***, p < 0.01 **, p < 0.05 *, p > 0.05 ^ns^.

### Benzoxazinoid accumulation co-varies with chemical soil gradient

To characterize the conditioning phase (**Fig. 1**), we collected soil samples at maize harvest for benzoxazinoid analysis. We confirmed the presence of benzoxazinoids in soils of wild-type plots while we did not detect them in plots where mutant *bx1* plants were grown (**Fig. 2C**). We measured high amounts of HMBOA and HDMBOA-Glc followed by MBOA, DIMBOA-Glc and DIMBOA, and low amounts of AMPO in soils of wild-type plots. The benzoxazinoid measurements varied strongly across replicates. Soil levels of several benzoxazinoids, in particular DIMBOA, gradually increased along PC1 (**Fig. 2C, Fig. S4**), from the north to the south of the field. Thus, the innate differences in soil chemistry are associated with marked changes in benzoxazinoid accumulation.

### Microbial community composition co-varies with chemical soil gradients

We also profiled the microbiota to describe the microbial communities of maize roots, its rhizospheres, and the soil at maize harvest. Again, we noticed that the variation in microbiota composition coincided with the position of the plot in the field along PC1 (**Fig. 2D**). Permutational multivariate analysis of variance (PERMANOVA) revealed significant positional effects for the bacteria and the fungi. Taking R^2^ values as indicators for effect size, positional effects on bacteria were stronger. The strong positional effects in bacterial communities were apparent in Principal Coordinate Analysis (PCoA), where the second axis largely separated the replicates following their position along PC1 (**Fig. S5**). Thus, the innate differences in soil chemistry are associated with differential microbial community composition.

### Benzoxazinoid exudation shapes root microbiota

To determine whether benzoxazinoids shape microbial communities in maize roots and rhizospheres, as observed before (Cadot *et al.*, 2021b), we first analysed, taking the soil chemical gradient into account, the impact of benzoxazinoids on alpha diversity. Alpha diversity of root and rhizosphere bacteria as well as rhizosphere fungi were enhanced in wild-type samples relative to *bx1* samples (**Fig. S6**). We then measured changes in beta diversity using PERMANOVA to validate benzoxazinoid conditioning and compare the effect size (R^2^ values) relative to PC1. Benzoxazinoid exudation shaped microbial communities in the roots and rhizospheres, with stronger effects of soil chemistry than benzoxazinoid effects (**Table S1**). Constrained Analysis of Principal Coordinates (CAP) visually confirmed the effects of benzoxazinoids and soil chemistry on microbial community composition (**Fig. 3**). Thus, benzoxazinoid exudation led to a microbial conditioning.

**Figure 3.**
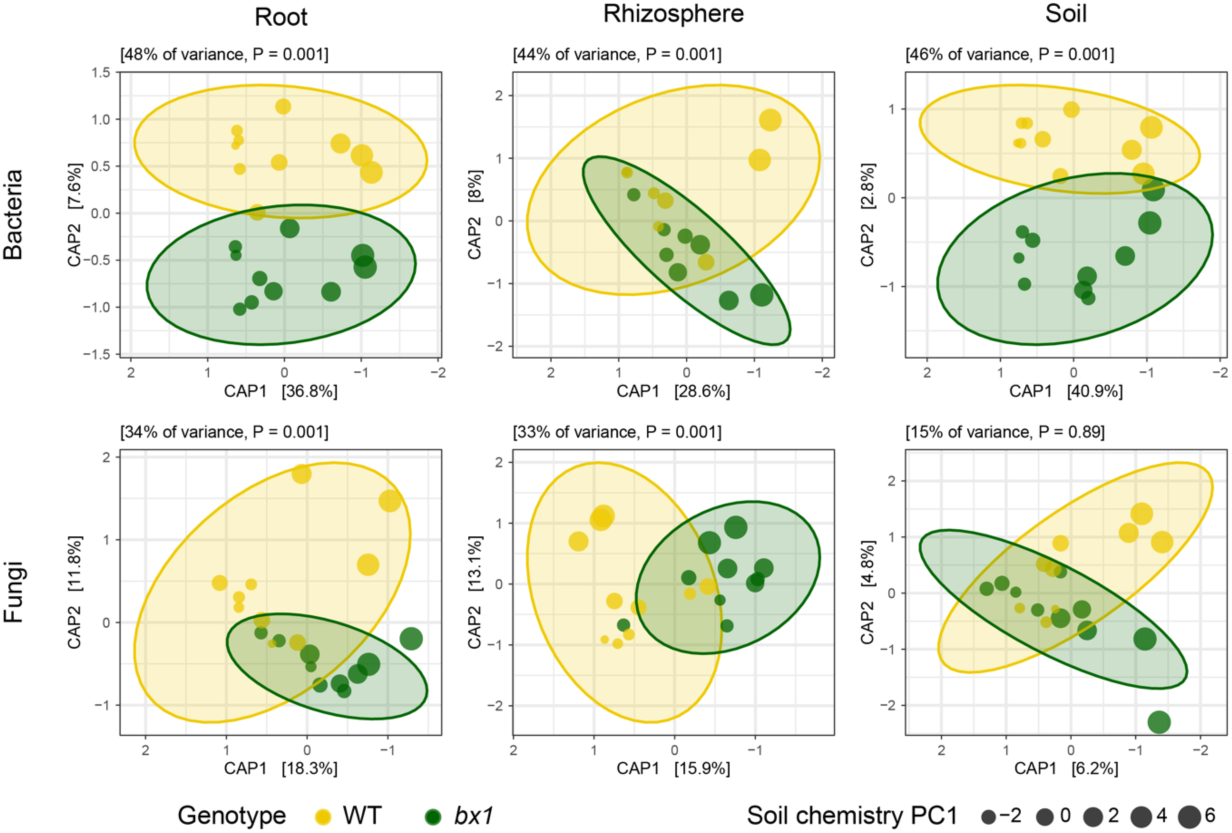
Benzoxazinoid-dependent structuring of rhizosphere and root microbiota. Compartment-wise Constrained Analysis of Principal Coordinates (CAP) using Bray-Curtis distances of community profiles from bacteria (top) and fungi (bottom). CAPs were performed using the model ‘∼ maize genotype * soil chemistry PC1’. Wild-type (WT) and *bx1* mutant samples are shown for roots, rhizospheres, and soils. Datapoints were sized using the values of soil chemistry PC1. Total variance explained by the model and model significance are shown at the top of each panel. Axis labels indicate percentage of variance explained.

### Chemical legacy of benzoxazinoid exudation persists in soil

To test the persistence of benzoxazinoid-dependent effects, we measured benzoxazinoid contents again at wheat sowing and during wheat growth in the feedback phase (**Fig. 1**). At wheat sowing we found 10-100 fold reduced levels of benzoxazinoids compared to our first measurements (**Fig. 4A**). We thus performed analyses on concentrated samples, which also resulted in the detection of low benzoxazinoid levels in *bx1* conditioned soils. Most benzoxazinoids were still significantly more abundant in soils of wild-type plots (**Fig. 4B**). These quantitative differences were lost during wheat vegetative growth and some compounds (HDMBOA-Glc, DIMBOA, and HMBOA) became more abundant compared to wheat sowing, as wheat also releases benzoxazinoids to the soil (**Fig. 4B**). AMPO, the microbial metabolization product of MBOA, behaved differently than the other benzoxazinoids (**Fig. 4A**): Its concentration decreased only marginally across time points, and it remained significantly higher in wild-type conditioned plots during the entire experiment (**Fig. 4B**). We observed a significant co-variation of AMPO with PC1 (**Fig. S7**). The concentration gradient of AMPO was always opposite to other benzoxazinoids such as HDMBOA-Glc, DIMBOA and HMBOA (**Fig. S4, Fig. S7**), suggesting that it may result from differential conversion of benzoxazinoids to their breakdown products.

**Figure 4.**
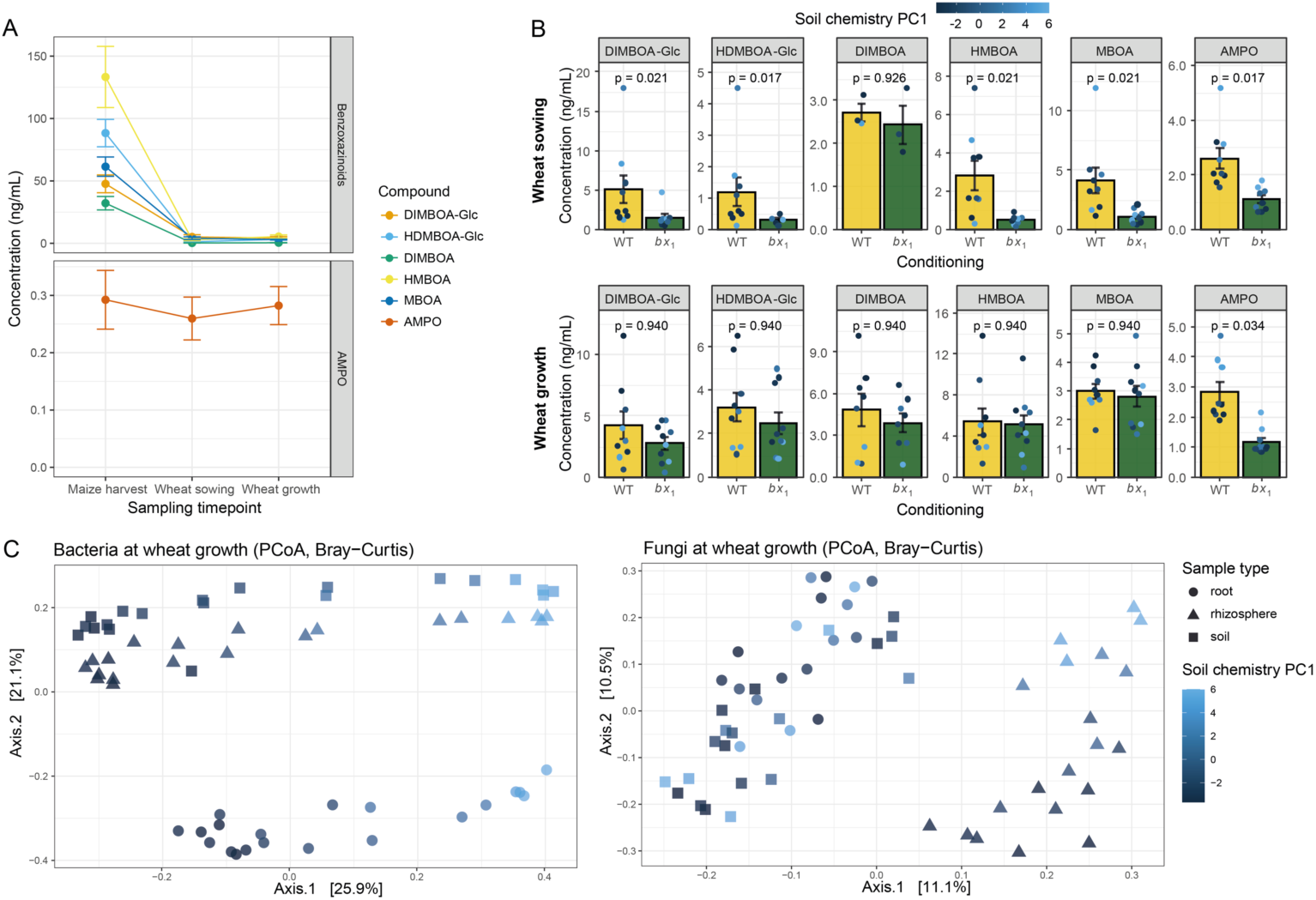
Benzoxazinoid-mediated chemical legacy persists in soil. (A) Progression of concentrations of benzoxazinoids and their degradation product (AMPO) in wild-type plots over time (Means ± SE). (B) Concentrations of benzoxazinoids in soils collected on plots conditioned by wild-type (WT) or benzoxazinoid-deficient *bx1* mutant plants in ng/mL of soil (Means ± SE) at wheat sowing (top) and during wheat growth (bottom). *P* values were calculated by Wilcoxon rank-sum tests and corrected for multiple testing. (C) Unconstrained Principal Coordinate Analysis (PCoA) using Bray-Curtis distances of bacterial (left) and fungal (right) communities in root, rhizosphere, and soil samples. Axis labels indicate percentage of variance explained.

To test the persistence of the microbial legacy found at maize harvest (**Fig. 3**), we profiled the soil microbiomes at wheat sowing, and the soil, rhizosphere, and root microbiomes again during wheat growth. PERMANOVA revealed significant effect sizes for the soil chemical gradient, at wheat sowing and during wheat growth (**Table S2**). Unconstrained PCoA visualized the structuring of the bacterial communities and of rhizosphere fungi by the soil chemical gradient (**Fig. 4C**). However, no significant impact of benzoxazinoid conditioning on the soil and wheat microbial community composition was detected (**Table S2**). Thus, while chemical legacies of benzoxazinoid exudation remained present during wheat growth, microbial legacies disappeared in the feedback phase.

### Soil chemistry directly determines benzoxazinoid degradation

Differences in soil chemistry may change benzoxazinoid exudation and/or their degradation, thus accounting for the marked gradient in the directionality of the observed benzoxazinoid accumulation and microbial community composition. To test this hypothesis, we sampled soil from the opposite ends of the soil chemical gradient, i.e. south (S) and north (N) soils (**Fig. 5A**). We then grew wild-type maize plants in these soils for 3 weeks and measured benzoxazinoid accumulation in the soil and benzoxazinoid exudation from freshly harvested roots. We did not detect significant differences in benzoxazinoid exudation from roots (**Fig. 5B**). However, we found significantly higher benzoxazinoid levels in S soil compared to N soil (**Fig. 5C**). The glycosylated benzoxazinoids and their conversion products were more abundant in soil of the S compared to the N corner; a finding consistent with the field measurements (**Fig. S4, Fig. S7**).

**Figure 5.**
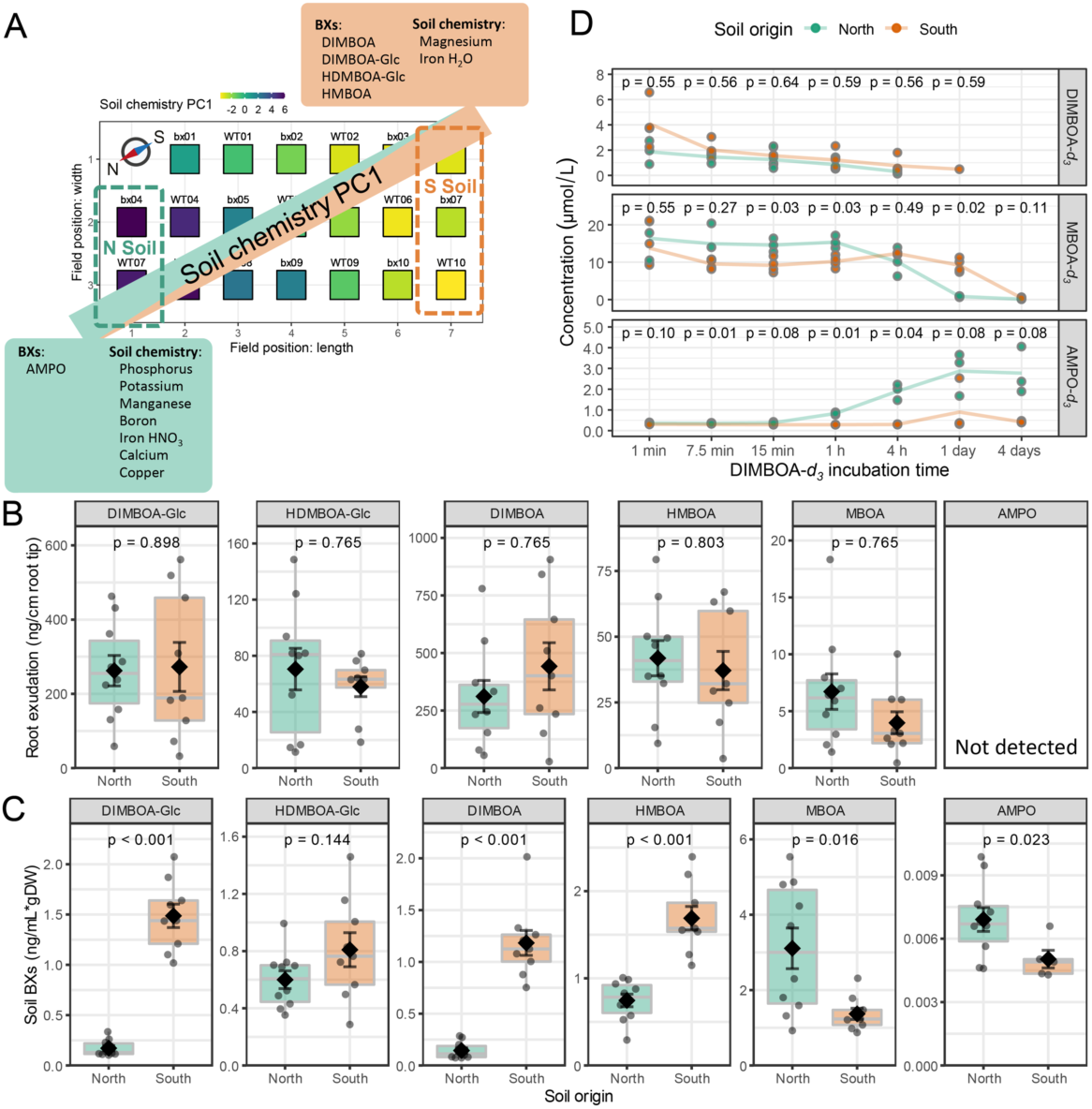
Soil chemistry-dependent degradation of benzoxazinoids. (A) Field soils at both extremes of the soil chemistry gradient were collected for benzoxazinoid exudation and degradation experiments under controlled conditions. Enriched chemical parameters in the north and south corners are listed in the mint and orange boxes, respectively (B) Root benzoxazinoid exudation of 3-week-old maize plants (W22). (C) Benzoxazinoid concentration in soils of 3-week-old maize plants (W22) measured in ng/mL of soil and corrected for root dry weight. (D) Degradation of deuterated DIMBOA-*d_3_* in a plant free system monitored for 4 days. For (B) and (C) boxplots, means ± SE, and individual datapoints are shown and for (B)-(D) outputs of Welch’s two-sample t-tests are included (FDR-corrected *p* values). N: north, S: south.

To further investigate benzoxazinoid metabolization, we performed an incubation experiment with labelled DIMBOA-*d_3_* directly spiked in S and N soils and quantified the benzoxazinoid degradation over time. Most of the DIMBOA was rapidly metabolized in the field soils (**Fig. S8**). In N soil, DIMBOA was metabolized to MBOA more rapidly, resulting in a faster and stronger accumulation of AMPO compared to S soil (**Fig. 5D**). In the S soil, almost no AMPO was formed despite complete metabolization of DIMBOA and MBOA, suggesting that other degradation pathways operate in this corner of the field. Overall, these experiments revealed that benzoxazinoid metabolization is strongly dependent on soil properties, which explains the strong gradient of benzoxazinoids and their degradation products observed across the different plots of the field experiment.

### Chemical soil gradients are associated with benzoxazinoid-dependent plant-soil feedbacks

To determine the effect of maize benzoxazinoid soil conditioning on the following crop along the soil chemical gradient, we measured wheat performance and resistance in the different plots. For each phenotype, we tested for benzoxazinoid-dependent feedback effects, effects of the soil chemical gradient (PC1) and their interaction. We also quantified the *local feedback* for each plot individually as log-response ratio of wild-type relative to *bx1* soil conditioning at a given location (see **Fig. S1**). This approach allowed us to compute local benzoxazinoid effects.

Overall, seedling emergence was not affected by soil conditioning or PC1 (**Fig. 6A**). Analysis of local effects however revealed a negative effect of benzoxazinoids in plots to the north, and a positive effect in the plots to the south.

**Figure 6.**
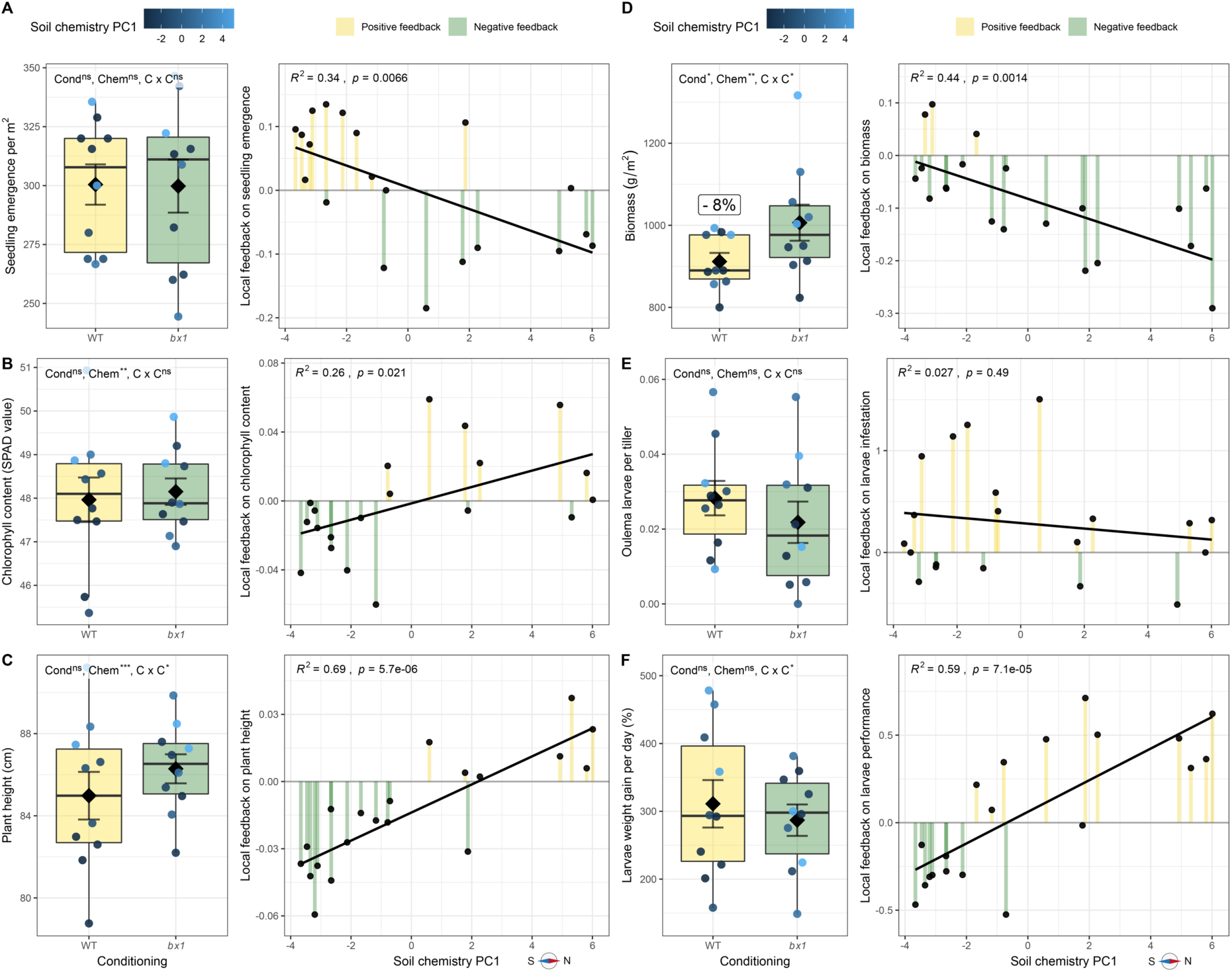
Benzoxazinoid-dependent feedbacks during wheat emergence and growth are associated with soil chemistry. (A) Seedling emergence, (B) chlorophyll content, (C) plant height, (D) dry biomass, (E) *O. melanopus* infestation, and (F) *S. littoralis* performance during wheat growth. For each phenotype boxplots (left) and local feedbacks of individual plots along the soil chemistry PC1 (right) are shown. For boxplots, phenotypes measured on plots conditioned by wild-type (WT) or benzoxazinoid-deficient *bx1* mutant maize are shown. Means ± SE and individual datapoints are included. Further, significance of ANOVA output is shown, where benzoxazinoid soil conditioning (Cond), the soil chemistry PC1 (Chem), and their interaction (C x C) were modelled. For the local feedbacks, log-response ratio values of individual plots are shown and *R^2^* and *p* value of linear regression are indicated on top. For more details on the local feedback refer to method section. Levels of significance: p < 0.001 ***, p < 0.01 **, p < 0.05 *, p > 0.05 ^ns^.

During wheat growth, overall chlorophyll content and height were not affected by benzoxazinoid soil conditioning, but local effects were again detected (**Fig. 6B-C**). Positive effects of benzoxazinoids on chlorophyll and height were observed in plots to the north, while negative effects were observed in plots to the south. Plant biomass was negatively affected by benzoxazinoid soil conditioning, with effects that were more pronounced towards the northern end of the gradient in the field (**Fig. 6D)**.

As defence-related phenotypes, we counted the number of *Oulema melanopus* larvae on the plants in the field and we tested the performance of *Spodoptera littoralis* feeding on leaf material collected in the field. Both defence phenotypes were not affected by benzoxazinoid soil conditioning (**Fig. 6E-F**). For *S. littoralis* performance, analysis of local effects revealed a positive effect of benzoxazinoids on larval growth in plots to the north, and a negative effect in the plots to the south.

At wheat harvest, no significant benzoxazinoid effects on shoot biomass, biomass per tiller, and tiller density were found (**Fig. 7A-C**) and the overall yield was also not affected by benzoxazinoid conditioning(**Fig. 7D**). However, for yield a weak effect was observed along the gradient, with positive effects in plots to the north and slightly negative effects in plots to the south.

**Figure 7.**
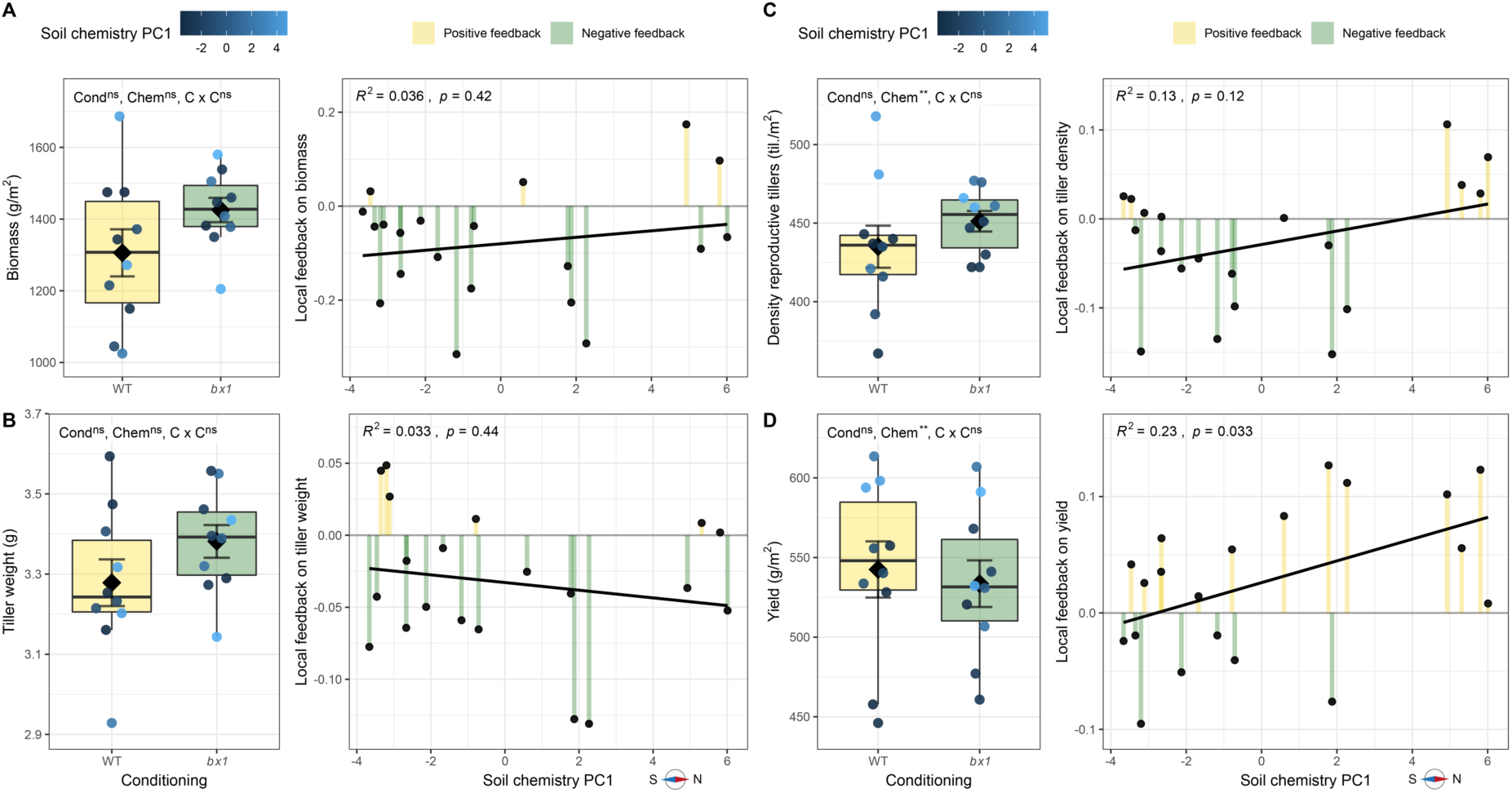
Local benzoxazinoid-dependent feedbacks on grain yield at wheat harvest. (A) Biomass, (B) tiller weight, (C) tiller density, and (D) yield at wheat harvest. For each phenotype, boxplots (left) and local feedbacks of individual plots along the soil chemistry PC1 (right) are shown. For boxplots, phenotypes measured on plots conditioned by wild-type (WT) or benzoxazinoid-deficient *bx1* mutant maize are shown. Means ± SE and individual datapoints are included. Further, significance of ANOVA output is shown, where benzoxazinoid soil conditioning (Cond), the soil chemistry PC1 (Chem), and their interaction (C x C) were modelled. For the local feedbacks, log-response ratio values of individual plots are shown and *R^2^* and *p* value of linear regression are indicated on top. For more details on the local feedback refer to method section. Levels of significance: p < 0.001 ***, p < 0.01 **, p < 0.05 *, p > 0.05 ^ns^.

Agronomically important kernel quality parameters including grain characteristics, protein content, and bakeability were all not affected by overall benzoxazinoid soil conditioning (**Fig. S9**). Gradients of feedback effects on grain width, volume weight and dough stability were detected. Nutritional and food quality properties were not changed by benzoxazinoid conditioning or along the soil chemical gradient (**Fig. S10**). Thus, benzoxazinoid soil conditioning influences wheat growth, defence, yield, and grain quality, but the directionality of the effect follows environmental gradients associated with differences in soil chemistry.

## Discussion

Soil conditioning by plant secondary metabolites can affect the growth and defence of the following crop through plant-soil feedbacks (Hu *et al.*, 2018b). If and how such feedbacks depend on the soil type, and how soil heterogeneity may influence their spatial patterning, remains unknown. Here, we show that the effect of maize benzoxazinoids on wheat performance is was fully dependent on soil properties, leading to a distinct effect gradient within a single field. Correlation analysis revealed strong associations between soil parameters (chemistry, microbiome), soil benzoxazinoid accumulation, and the magnitude and direction of the benzoxazinoid-dependent feedback effects on wheat growth, defence, and grain quality. Below, we discuss these findings from a mechanistic perspective and derive implications for the use of secondary metabolite-driven plant-soil feedbacks in agriculture.

### Impact of soil chemistry on benzoxazinoid accumulation

We find that innate differences in soil chemistry are associated with marked variation in benzoxazinoid accumulation in soil (**Figs. 2C, S4, 4B, S7**). Benzoxazinoid exudation can, for example, be altered in response to soil iron (Zhou *et al.*, 2018) and aluminium (Zhao *et al.*, 2019). Soil parameters may also influence the metabolization of secondary metabolites (Nannipieri *et al.*, 2002). The degradation to MBOA for instance is pH dependent (Maresh *et al.*, 2006), and the conversion of MBOA to AMPO as well as AMPO metabolization are mediated by soil microbes (Etzerodt *et al.*, 2008; Niemeyer, 2009). Our climate chamber experiments show that the differential accumulation in soils from different field positions is the result of differences in metabolization rather than exudation by maize roots (**Fig. 5**). As benzoxazinoids in the rhizosphere are directly responsible for changes in microbial composition and feedback effects on other plants (Hu *et al.*, 2018b), differences in metabolization may influence plant-soil feedback. The pronounced differences in metabolization we observe within the same field point to substantial potential for fine-scale variation in secondary metabolite-mediated feedback effects.

### Interactions between benzoxazinoids and microbiota

Root-associated bacterial and fungal community compositions are well documented to be affected by benzoxazinoid exudation (Hu *et al.*, 2018b; Cotton *et al.*, 2019; Kudjordjie *et al.*, 2019; Cadot *et al.*, 2021b; Gfeller *et al.*, 2022). Here, we confirm this result and show that benzoxazinoid effects on microbiome composition and alpha diversity are significant, even in heterogeneous soils (**Fig. 3**, **Table S1**). The community structure of fungi, compared to bacteria, was more strongly affected by benzoxazinoids, which is in line with previous findings (Cadot *et al.*, 2021b). Bacterial communities showed a strong association with soil chemistry, possibly because they respond more dynamically to local changes in environmental conditions. Previous work showed that bacteria are, for example, more strongly affected by soil acidification compared to fungi (Rousk *et al.*, 2010; Choma *et al.*, 2020). Given that benzoxazinoid accumulation is dependent on variation in soil properties and the root and rhizosphere microbiota are shaped by benzoxazinoids, one would expect the benzoxazinoid effect on microbiomes to vary across the field. In our study we did not observe this behaviour. A possible explanation for this is that root and rhizosphere microbiota are shaped directly by root-exuded benzoxazinoids, which were shown to be unaffected by soil parameters (**Fig. 5B**). In line with our previous field study (Gfeller *et al.*, 2022), chemical, but not microbiota patterns persisted to the next crop generation. This is in contrast to previous pot and container experiments, where microbial fingerprints from the soil conditioning phase were still present during the feedback plant’s growth (Hu *et al.*, 2018b; Hannula *et al.*, 2021). A likely explanation for this discrepancy is that in our experiments, the process of seedbed preparation for wheat with a complete soil homogenization at a depth of 10 cm, resulted in the dilution of the microbial fingerprints in the surrounding soil. In summary, both soil chemistry and benzoxazinoid exudation shapes root microbiota, which likely adds to the variation and dynamics of plant-soil feedback effects.

### Within-field variation in plant-soil feedbacks

Plant-soil feedbacks are well known to depend on the environmental context, the responsible mechanisms are however only partly understood (van der Putten *et al.*, 2013; Smith-Ramesh & Reynolds, 2017; Gfeller *et al.*, 2023). Plant nutrient supply, for example, can influence the outcome of plant-soil feedbacks in crops and wild plants (Kos *et al.*, 2015a; Kuerban *et al.*, 2022). Generally, it is assumed that increasing soil fertility will weaken the strength of plant-soil feedbacks by lowering soil nutrient feedbacks. This would reduce the plant’s dependency on mutualists, and decrease the role of pathogens if plants have more resources to allocate in defence and immunity (Smith-Ramesh & Reynolds, 2017). Here, we found that depending on soil chemistry, the effect of benzoxazinoid-dependent plant-soil feedbacks on growth, defence, and food quality differ in strength and/or direction (**Figs. 6, 7**). During vegetative growth we found that under more fertile conditions, as indicated by higher wheat yield, benzoxazinoid conditioning led to faster plant growth, but less biomass accumulation and lower plant defence. Soil fertility positively correlated with benzoxazinoid degradation and affected microbial community composition. The exact underlying mechanism, however, remains to be investigated. The observed context dependency of plant-soil feedbacks within one field could explain why greenhouse experiments often cannot be reproduced under natural conditions (Schittko *et al.*, 2016; Forero *et al.*, 2019). We further found context dependencies of benzoxazinoid-dependent plant-soil feedbacks between studies: In this field experiment, at the end of wheat vegetative growth, we found an overall negative effect of benzoxazinoid conditioning on wheat biomass accumulation. This finding is in line with the conclusion of a previous greenhouse experiment (Cadot *et al.*, 2021a), but different from a previous field experiment (Gfeller *et al.*, 2022). Given that the two field experiments were conducted in different soils at different locations, the observed variation could be explained by benzoxazinoid-dependent plant-soil feedbacks being soil specific, as it was shown in this study and in a maize-maize experiment before (Cadot *et al.*, 2021a). Taken together, our findings show the importance to take into account local variation of plant-soil feedbacks to understand them in diverse environments and further examine their potential in sustainable agriculture.

### Conclusion

Plants closely interact with their belowground environment. Root exuded secondary metabolites can directly or indirectly, mediated through changes in the microbiome, affect the next plant generation grown in that soil. Our work shows that such secondary metabolite-mediated plant-soil feedbacks occur within crop rotations under agronomically relevant conditions and that they are highly dependent on the soil chemical context. Consistent with previous work showing that direct effects of benzoxazinoids on aboveground insects depend on soil chemistry (Hu *et al.*, 2021), this study highlights the importance of the local environmental context to drive the effects of plant secondary metabolites. The ultimate and resulting implication for agriculture is the necessity to understand the soil chemical and microbial context dependency of plant-soil feedbacks in crop rotations to make them applicable as a predictable and sustainable practice.

## Data availability

The raw microbiome sequencing data is available from the European Nucleotide Archive (http://www.ebi.ac.uk/ena) under the study accession PRJEB59165 and the sample IDs ERS14468209 and ERS14468210. All other data generated for this study and the R code to reproduce the statistical analyses are deposited on GitHub (https://github.com/PMI-Basel/Gfeller_et_al_Changins_field_experiment).

## Acknowledgements

We specifically thank Nicolas Widmer (Merci!) and Lilia Levy from Agroscope Changins (Switzerland) for access and assistance during the two-year field experiment. Further, we thank Florian Enz for field and laboratory assistance. This work was supported by the Swiss State Secretariat for Education, Research and Innovation (grant C15.0111 to KS and ME) and the Interfaculty Research Collaboration “One Health” of the University of Bern.

## Supporting Information

## Supplementary Methods

### Field experiment

The field experiment was conducted in Changins (Nyon, Switzerland) on a field at Agroscope (Parcel 29, 46°23’58.6”N, 6°14’23.1"E; **Fig. 1**) in the growing seasons of 2018 and 2019. The field was first ploughed (20 cm depth) on April 24^th^, and then harrowed (10 cm depth) on the 25^th^. On April 26^th^ the maize seeds were sown (200-220 seeds for a 20 m^2^ plot surface) on 20 plots (10 plots with wild-type plants alternating with 10 plots with *bx1* mutants). Each plot was 6 m long and 3 m wide and separated from each other by 3 m of buffer (**Fig. 1**). The six maize rows sown per plot were separated by 50 cm, and plants in the rows were separated by 20 cm. A net (14 g/m^2^) was put on the field during germination to protect seeds from being eaten by birds. On May 26^th^, alfalfa (*Medicago sativa*) was sown as a buffer plant between the plots and around the experimental field. The same day, ammonium nitrate (27.5%, 100 kg/ha) was used as N fertilizer and the soil was superficially weeded incorporating the fertilizer. On July 24^th^, 35 mm water was applied on the field for irrigation. No herbicides were utilized but manual weeding was performed on June 12^th^. The preceding crops were alfalfa (*Medicago sativa*, 2016-2017), spring wheat (2015), and maize (2014). On September 6^th^, the maize plants were harvested at maturity.

### Kernel analysis

To determine morphological kernels traits, an aliquot of 25 mL was taken. By means of a MARVIN kernel analyzer (GTA Sensorik GmbH, Germany) and a microbalance, volume weight, thousand kernel weight (TKW), kernel surface area, kernel length, and kernel width were determined. To test flour quality, an aliquot of kernels was milled and falling number (according to ICC standard method 107/1), Zeleny index (according to ICC standard method 116/1), and protein content were analysed, where protein content was evaluated by near-infrared reflectance spectroscopy (NIRS) using a NIRFlex N-500 (Büchi Labortechnik AG, Switzerland). Further, dough quality was determined using the micro-doughLAB farinograph (model 1800, Perten Instruments, PerkinElmer United States), by measuring dough stability (min), dough softening (Farinograph Units, FU), and water absorption capacity of the flour (%) during the kneading process according to the manufacturer’s protocol.

To test for possible benzoxazinoid conditioning effects on food quality related parameters, we send 750 g of kernels per plot to Eurofins Scientific AG (Schönenwerd, Switzerland) to analyse the most important nutritional values and mycotoxins.

### Benzoxazinoid analysis

To determine the dynamics of benzoxazinoids and their degradation products in soil over the course of the experiment, we analysed soil benzoxazinoid concentrations at the end of maize soil conditioning, after wheat sowing, and at the end of wheat vegetative growth. Samples were collected as described above, soils were passed through a test sieve (5 mm mesh size), 25 mL of soil was transferred into a 50 mL centrifuge tube and completely suspended in 25 mL acidified MeOH/H_2_O (70:30 v/v; 0.1% formic acid) by vigorously shaking and vortexing the tube. After 30 minutes of shaking at room temperature in a rotary shaker, samples were centrifuged (5 min, 2’000 x g) to sediment the soil. The supernatant was passed through a filter paper (Grade 1; Size: 185 mm; Whatman, GE Healthcare Live Sciences), 1 mL of the flow through was transferred into a 1.5 mL centrifuge tube, centrifuged (10 min, 19’000 x g, 4 °C), and the supernatant was sterile filtered (Target2TM, Regenerated Cellulose Syringe Filters, pore size: 0.45 µm; Thermo Scientific) into a glass tube for further analysis.

All samples collected at wheat sowing and wheat vegetative growth were concentrated 20x before the second centrifugation step. For that, 20 mL of soil extract per sample was dried (45 °C; CentriVap, Labconco) and resuspended in 1 mL of acidified MeOH/H_2_O (70:30 v/v; 0.1% formic acid).

Benzoxazinoids and degradation products were analysed with an Acquity UHPLC system coupled to a G2-XS QTOF mass spectrometer (Waters AG, Bade-Dättwil, Switzerland) as previously described (Gfeller *et al*., 2022). Absolute quantification was done through standard curves of pure compounds. For that, MBOA (6-methoxy-benzoxazolin-2(3H)-one) was purchased from Sigma-Aldrich Chemie GmbH (Buchs, Switzerland). DIMBOA-Glc (2-O-β-D-glucopyranosyl-2,4-dihydroxy-7-methoxy-2H-1,4-benzoxazin-3(4H)-one) and HDMBOA-Glc (2-O-β-D-glucopyranosyl-2-hydroxy-4,7-dimethoxy-2H-1,4-benzoxazin-3(4H)-one) were isolated from maize plants in our laboratory. DIMBOA (2,4-dihydroxy-7-methoxy-2H-1,4-benzoxazin-3(4H)-one), DIMBOA-*d_3_* (2,4-dihydroxy-7-(methoxy-*d_3_*)-2H-1,4-benzoxazin-3(4H)-one), HMBOA (2-hydroxy-7-methoxy-2H-1,4-benzoxazin-3(4H)-one), and AMPO (9-methoxy-2-amino-3H-phenoxazin-3-one) were synthesized in our laboratory.

### Microbiota profiling

#### Sample preparation

For the microbiota profiling the samples were collected as described above followed by sample preparation as previously described (Gfeller *et al*., 2022). In short, soil samples were obtained by gently removing soil from the root systems and passing them through a 5 mm test sieve. Roots were cut at a soil depth of 5 cm to 15 cm, placed in a 50 mL centrifuge tube, and washed 4x with 25 mL of sterile deionized water by vigorously shaking them 10 times. Roots samples, containing endophytes and epiphytes, were then freeze-dried, and milled to fine powder using a Ball Mill (Retsch GmBH; 30 s at 30 Hz using one 1 cm steel ball). To prepare the rhizosphere samples, the first two washes of the root cleaning were combined, centrifuged (5 min at 3000 x g), and the resulting pellet was frozen at -80 °C before further processing. *DNA extraction:* DNA was extracted using the Spin Kit for Soil (MP Biomedical, USA), following the instructions of the manufacturer. For that, 20 mg of roots powder or 200 mg for rhizosphere or soil were taken. DNA concentrations were evaluated by means of an AccuClear Ultra High Sensitivity dsDNA Quantitation Kit (Biotium, USA).

#### PCR protocol

Bacterial and fungal community profiling were performed following a two-step PCR profiling protocol described in (Gfeller *et al*., 2022) with a few changes. Bacterial profiles are based on PCR primers 799-F (Chelius & Triplett, 2001) and 1193-R (Bodenhausen *et al*., 2013) that span the hypervariable regions V5 to V7 of the 16S rRNA gene. Fungal profiles are derived from the internal transcribed spacer region 1 (ITS1) and were amplified with the PCR primer pair ITS1-F (Gardes & Bruns, 1993) and ITS2 (White *et al*., 1990). The PCR reactions mix included 5-Prime HotMastermix (1x, QuantaBio, USA), bovine serum albumin BSA (0.3%), forward primer (300 nM), reverse primer (300 nM), with 1 ng input DNA in the case of soil and rhizosphere bacterial samples, 10 ng for root bacterial samples, 2 ng for soil and rhizosphere fungal samples and 20 ng for root fungal samples; water was added to the solution to obtain 20 uL. One negative control with no DNA and one validated positive control were added to each PCR plate. PCR settings for bacteria included a first 3-min step at 94 °C for denaturation, and 25 cycles of 45 s at 94 °C, 1 min at 50 °C, 1 min 30 s at 72 °C, followed by a final step of 10 min at 72 °C. The PCRs for fungal samples were done similarly, except that the number of cycles was increased from 25 to 35 to increase amplification. Amplification success and absences of contamination were confirmed by migrating PCR product aliquots on a 1.5% agarose gel. Next, PCR products were purified using self-made Solid Phase Reversible Immobilisation (SPRI) magnetic beads (https://openwetware.org/wiki/SPRI_bead_mix) with a 0.8:1 beads (16 µl) to PCR products (20 µl) ratio in 10 mM Tris-HCl pH 8 buffer solution. After binding the beads with the adhering DNA to a magnet, the supernatant was removed, the beads were washed twice with 80% ethanol, briefly air-dried, and eluted in 22.5 µl Tris-HCl buffer (10 mM pH 8). 20 µl of cleaned amplicon DNA were then transferred to new 96-well plates. These clean PCR products were then quantified with NanoDrop (Thermo Fisher, USA), equimolarly pooled, purified, and concentrated with beads in a 1:1 beads-to-library ratio, and eluted in 20 µl of buffer. Finally, the library was quantified with Qubit (Thermo Fisher, USA). Library preparation was completed by ligation of the Illumina adapters by the Next Generation Sequencing Platform at University of Bern, where the sequencing was performed on a Illumina MiSeq instrument (v3 chemistry, 300 bp paired end).

#### Bioinformatics

The sequencing data was processed as previously described (Gfeller *et al*., 2022). In short, raw reads were quality checked and demultiplexed using *FastQC* and *cutadapt*, respectively (Andrews, 2010; Martin, 2011). Exact amplicon sequences variants (ASVs) were generated using the *DADA2* pipeline (Callahan *et al*., 2016; R package *DADA2*). Taxonomic assignment of ASVs was performed using a *DADA2* formatted version of the *SILVA* v.132 database (Quast *et al*., 2013; Callahan, 2018) for bacteria and the FASTA general release from *UNITE* v8.1 (Nilsson *et al*., 2019) for fungi.

### Statistical analysis of microbiota profiles

Analyses were conducted using the open-source software R (R Core Team, 2021). Root, rhizosphere, and soil microbiomes were analysed at maize harvest, soil microbiomes at wheat sowing, and again all three compartments during wheat growth. ASVs were first filtered to exclude sequences assigned to eukaryotes, cyanobacteria, mitochondria, or chloroplasts. Based on the inspection of the sequencing depth of all samples and testing if sequencing depth was significantly different among variable groups by a Kruskal-Wallis test, microbial community data was rarefied for all downstream analysis using the v*egan* package (Weiss *et al*., 2017; Oksanen *et al*., 2020). Thresholds for rarefaction were 6701 for bacteria and 186 for fungi, resulting in the loss of four bacterial samples with sequence numbers below the threshold. A rarefaction curve revealed that samples were sequenced deep enough to cover microbial diversity.

First, we examined the microbial community composition along the soil chemical gradient (including PC1) after maize growth. Unconstrained Principal Coordinate Analysis (PCoA) using Bray-Curtis distances were performed for bacterial and fungal communities and the effect of compartment and soil chemistry on community composition were tested by permutational analysis of variance (PERMANOVA, 999 permutations) on Bray-Curtis distances using the R package *vegan* (Oksanen *et al*., 2020). The correlation between the soil chemistry (PC1) and microbial communities (PCoA axes) were further investigated by Pearson’s correlation tests. Next, we investigated the effect of benzoxazinoid exudation on alpha and beta diversities by comparing samples from wild-type and *bx1* plots within each compartment (root, rhizosphere, soil). Alpha diversity was analysed by calculating the Shannon index in each sample (based on the mean of 100 iterations) and performing an ANOVA, where we included soil chemistry (PC1) to account for otherwise unexplained variance (model: diversity ∼ genotype * soil chemistry PC1). The same model was used for PERMANOVA. We visualized beta diversity by plotting the Canonical Analysis of Principal coordinates (CAP) using the R package *phyloseq* (McMurdie & Holmes, 2013).

To test if the differences in microbial communities found at the end of maize growth persisted, we again analysed the soil microbiota at wheat sowing and soil, rhizosphere, and root microbiota during wheat growth. This was done by compartment-wise PERMANOVAs for bacteria and fungi at each sampling time and unconstrained PCoA visualization of the structuring microbiomes during wheat growth. All code for statistical analysis and visualization and the corresponding data can be downloaded from GitHub (https://github.com/PMI-Basel/Gfeller_et_al_Changins_field_experiment).

## Supplementary Figures

**Supplementary Figure S1.**
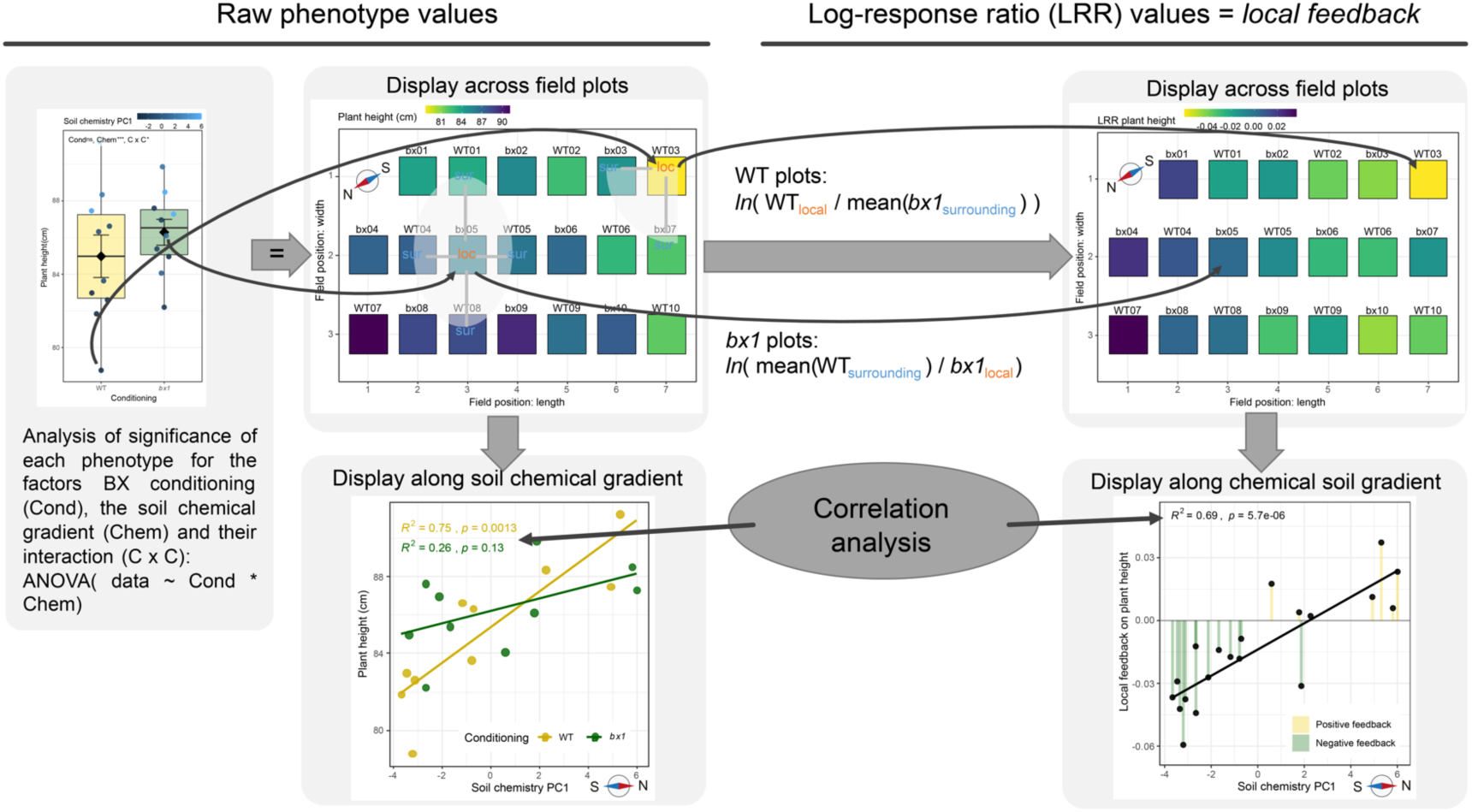
Explanation of the local feedback. Two different statistical analysis were performed on wheat phenotypes. First, the raw data was inspected for overall differences depending on benzoxazinoid soil conditioning and correlations with the soil chemical gradient. Second, the log-response ratio (LRR) was calculated for each plot, to estimate the local feedback at a given position on the field. To do so, on wild-type (WT) conditioned plots, the phenotype measurement was divided by the mean of measurements on the surrounding *bx1* mutant conditioned plots and log transformed (natural logarithm). On *bx1* conditioned plots, the mean phenotype measurement of the surrounding wild-type plots was divided by the measurement on the focal *bx1* plot and log transformed. These local feedbacks were further used to test for associations between soil parameters and the direction and strength of the feedback, where positive LRRs denote positive benzoxazinoid plant-soil feedbacks and negative LRRs denote negative benzoxazinoid-dependent plant-soil feedbacks.

**Supplementary Figure S2.**
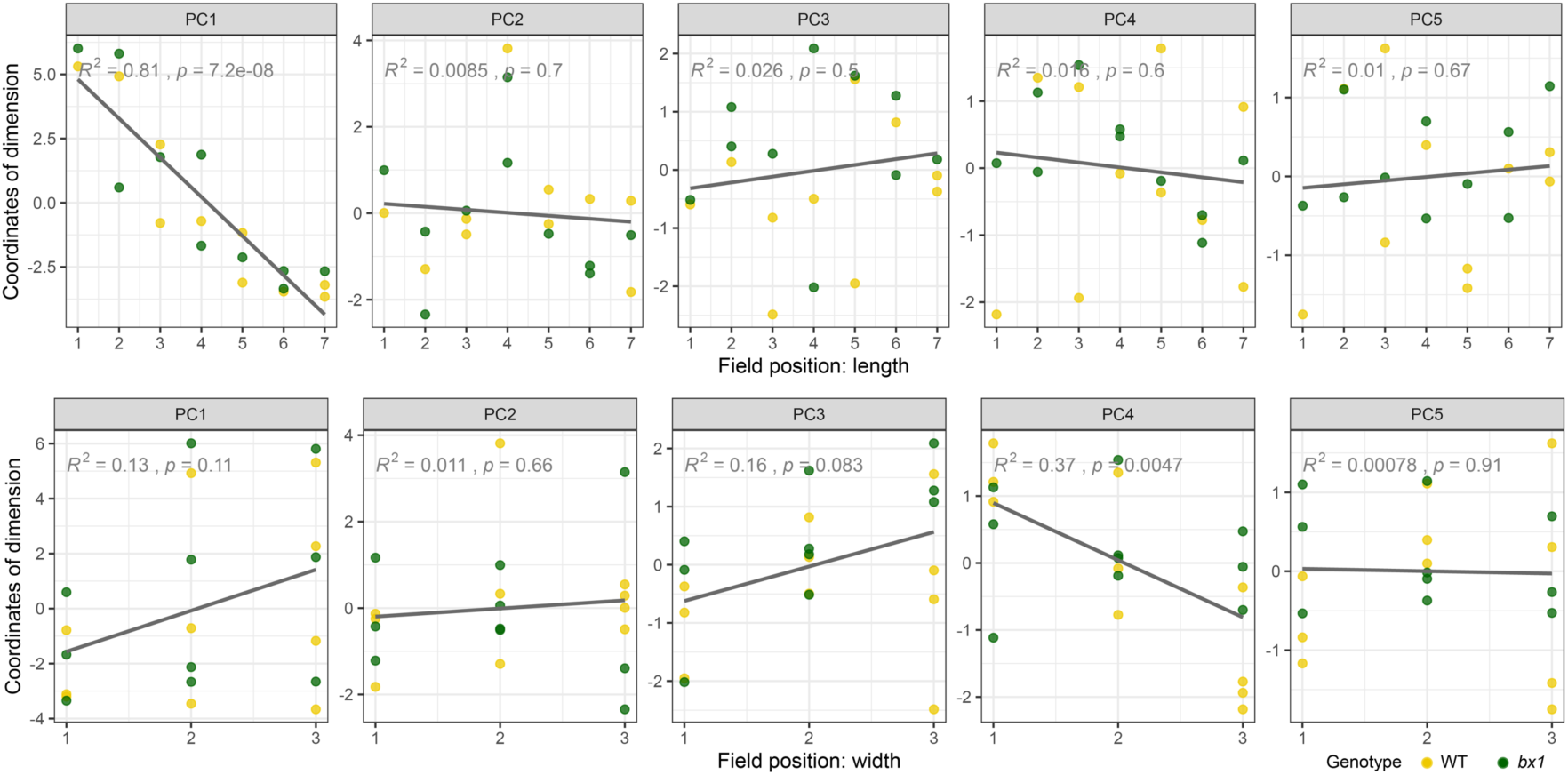
Composition of soil chemistry (PCA dimensions) varies with field positions. The correlation between soil chemistry Principal Component (PC) axes 1-5 and the position along field length (top) or field width (bottom) are shown. *R^2^* and *p* value of linear regression are indicated in the top. Yellow circles: wild-type (WT) plots; Green circles: *bx1* mutant plots.

**Supplementary Figure S3.**
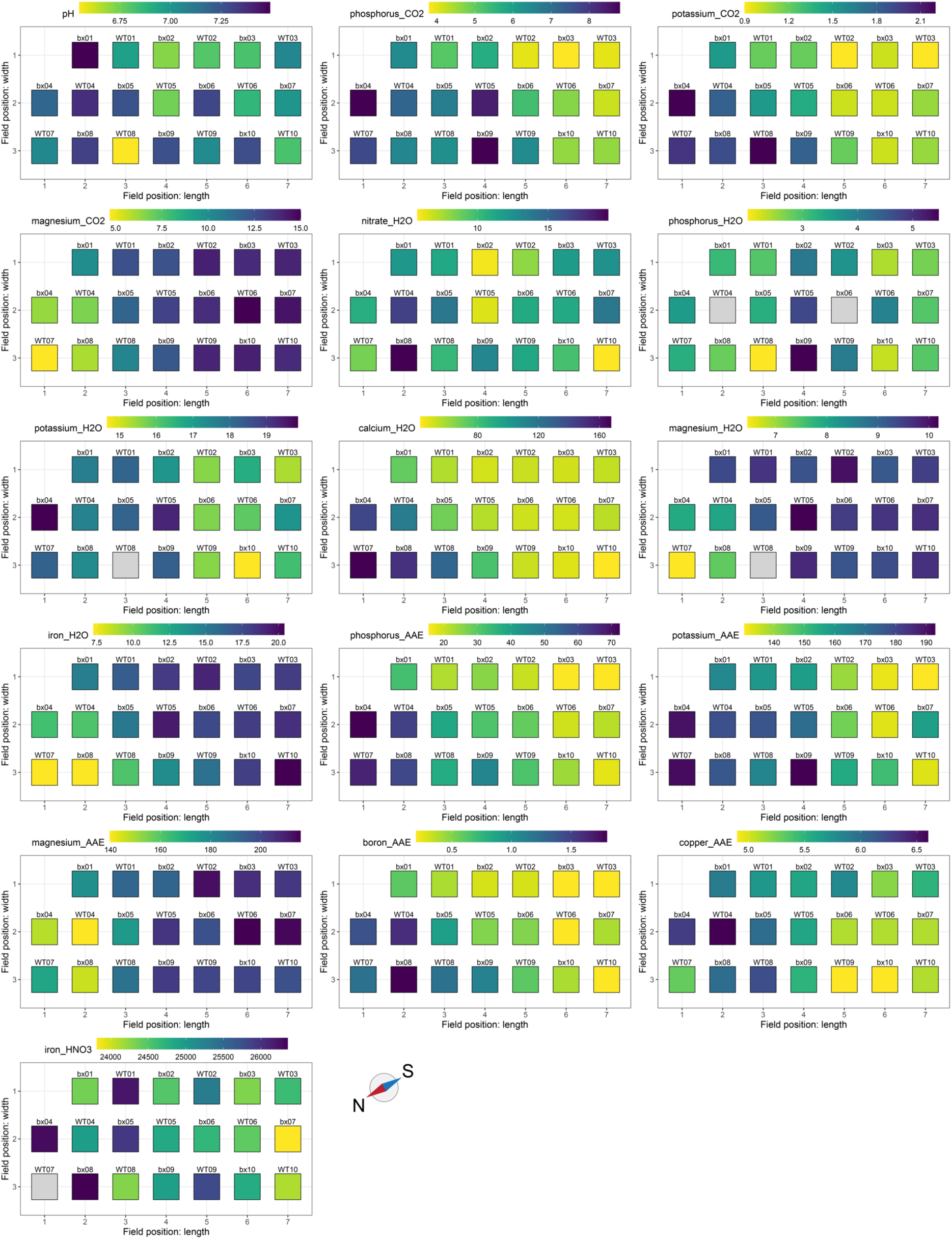
Chemical gradient along the field. Field map showing the soil chemical gradient along field plots for all individual soil parameters.

**Supplementary Figure S4.**
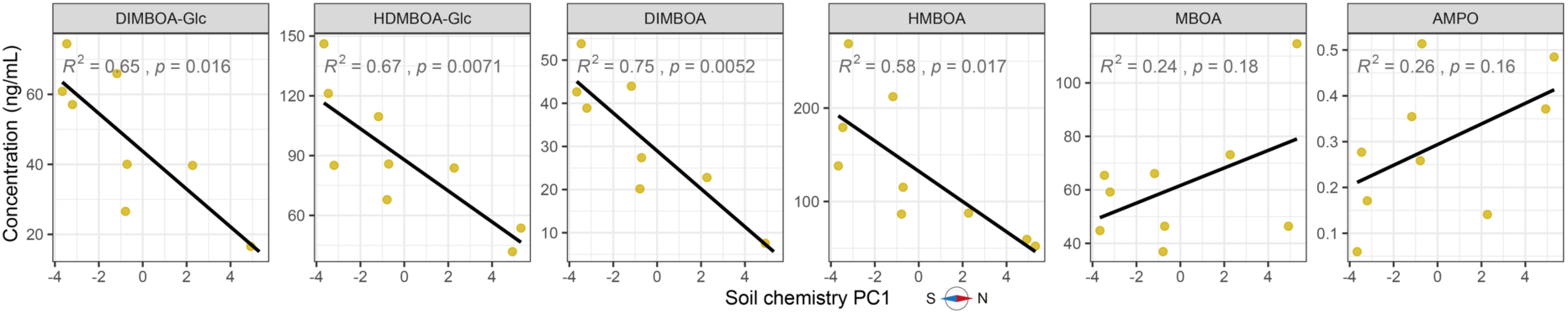
Soil benzoxazinoid concentrations at maize harvest are correlated with soil chemistry. The correlation between soil chemistry PC1 and the concentrations of benzoxazinoids in soils surrounding wild-type (WT) roots at the end of the condition phase in ng/mL of soil are shown. *R^2^* and *p* value of linear regression are in the top.

**Supplementary Figure S5.**
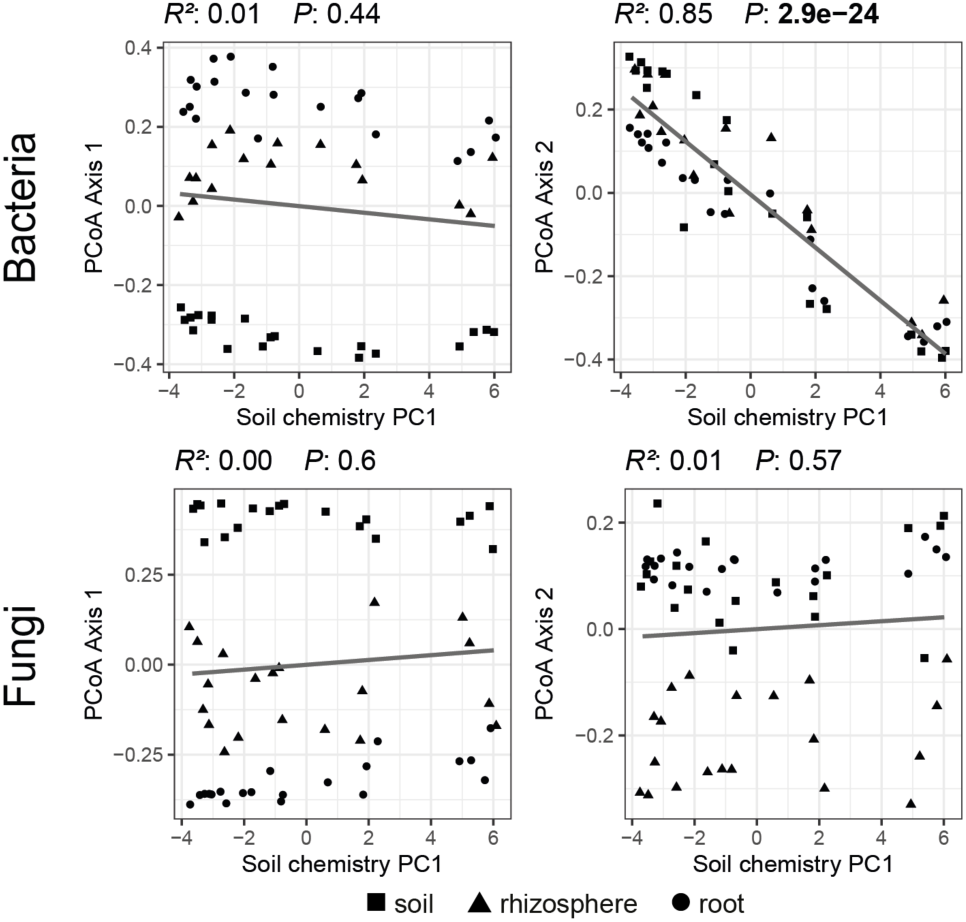
Microbial community composition (PCoA axes) varies with soil chemical gradient. Axis form Principal Coordinate Analysis (PCoA) of bacterial (top) and fungal (bottom) communities in roots, rhizospheres and soils samples. Pearson correlation and corresponding significance is denoted on top of each panel.

**Supplementary Figure S6.**
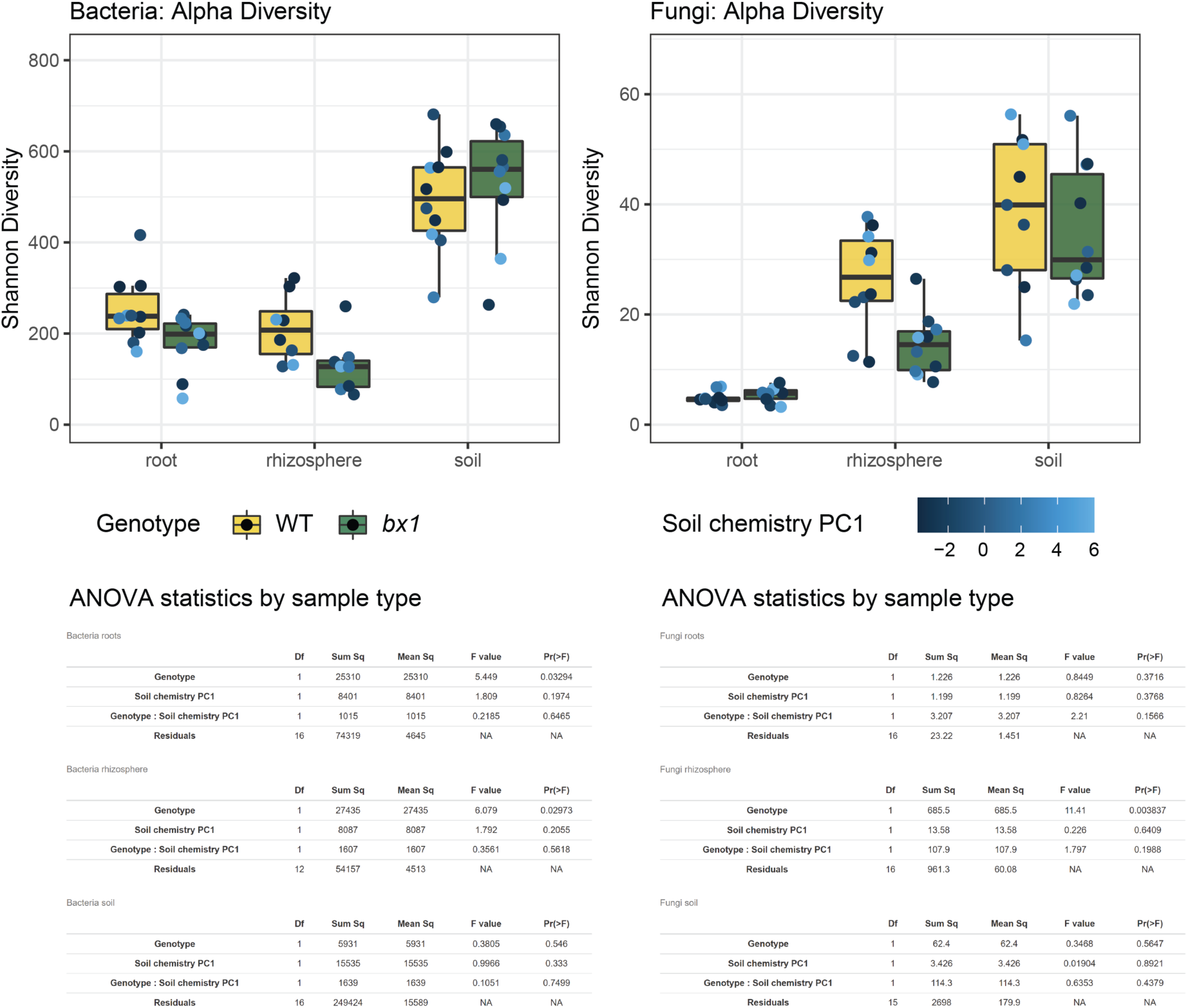
Microbial alpha diversity at maize harvest. (A) Bacterial and (B) fungal alpha diversity calculated as Shannon index are shown in boxplots (top) and compartment wise ANOVA outputs are included (bottom; model ‘∼ genotype * soil chemistry PC1’).

**Supplementary Figure S7.**
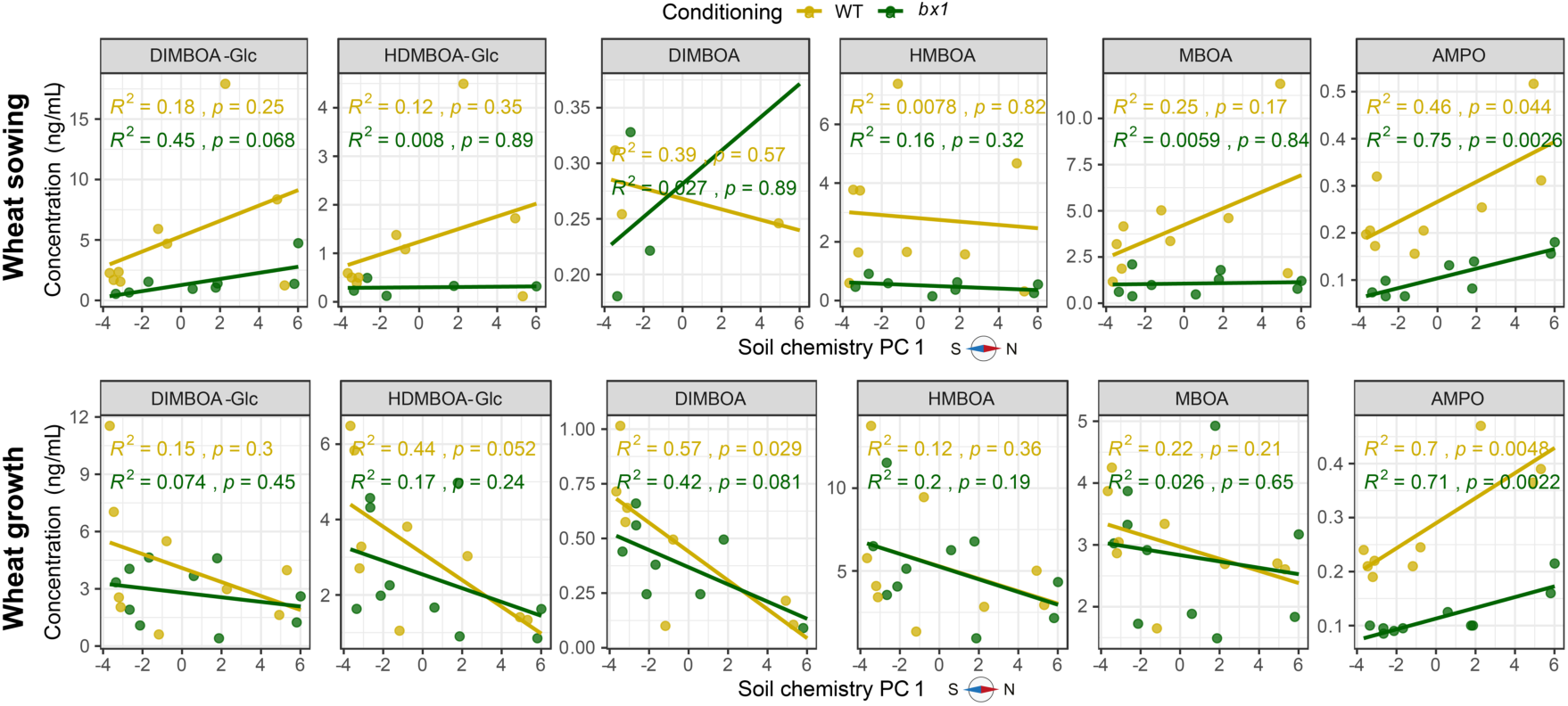
Soil benzoxazinoid measurements at wheat sowing and during wheat growth. The correlation between soil benzoxazinoid concentration and soil chemistry PC1 are shown at wheat sowing (A) and during wheat growth (B). *R^2^* and *p* values of linear regressions for plots conditioned by wild-type (WT) or benzoxazinoid-deficient *bx1* mutant plants are indicated in the top.

**Supplementary Figure S8.**
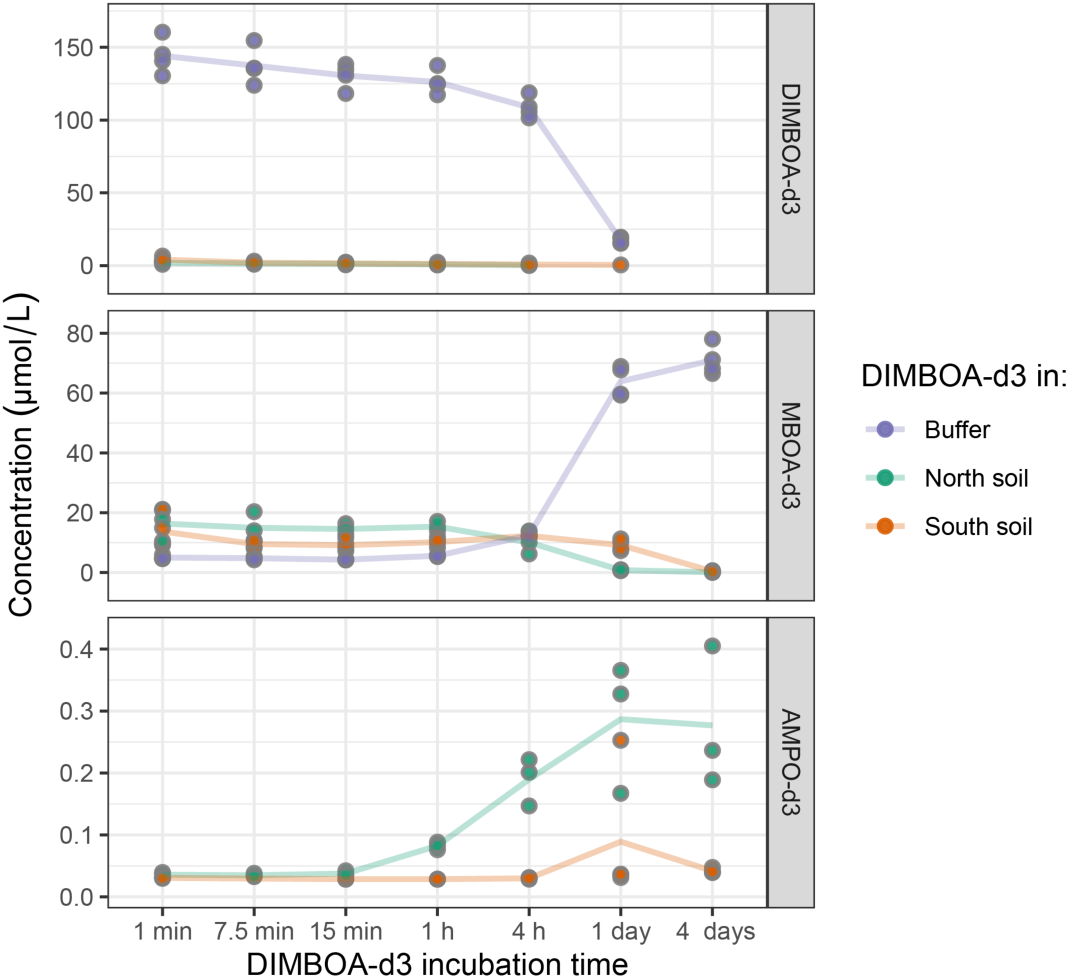
Degradation of benzoxazinoids in soil suspensions and buffer. Degradation of deuterated DIMBOA-*d_3_* in soil suspensions originating from both extremes of the soil chemistry gradient and in buffer was monitored for 4 days.

**Supplementary Figure S9.**
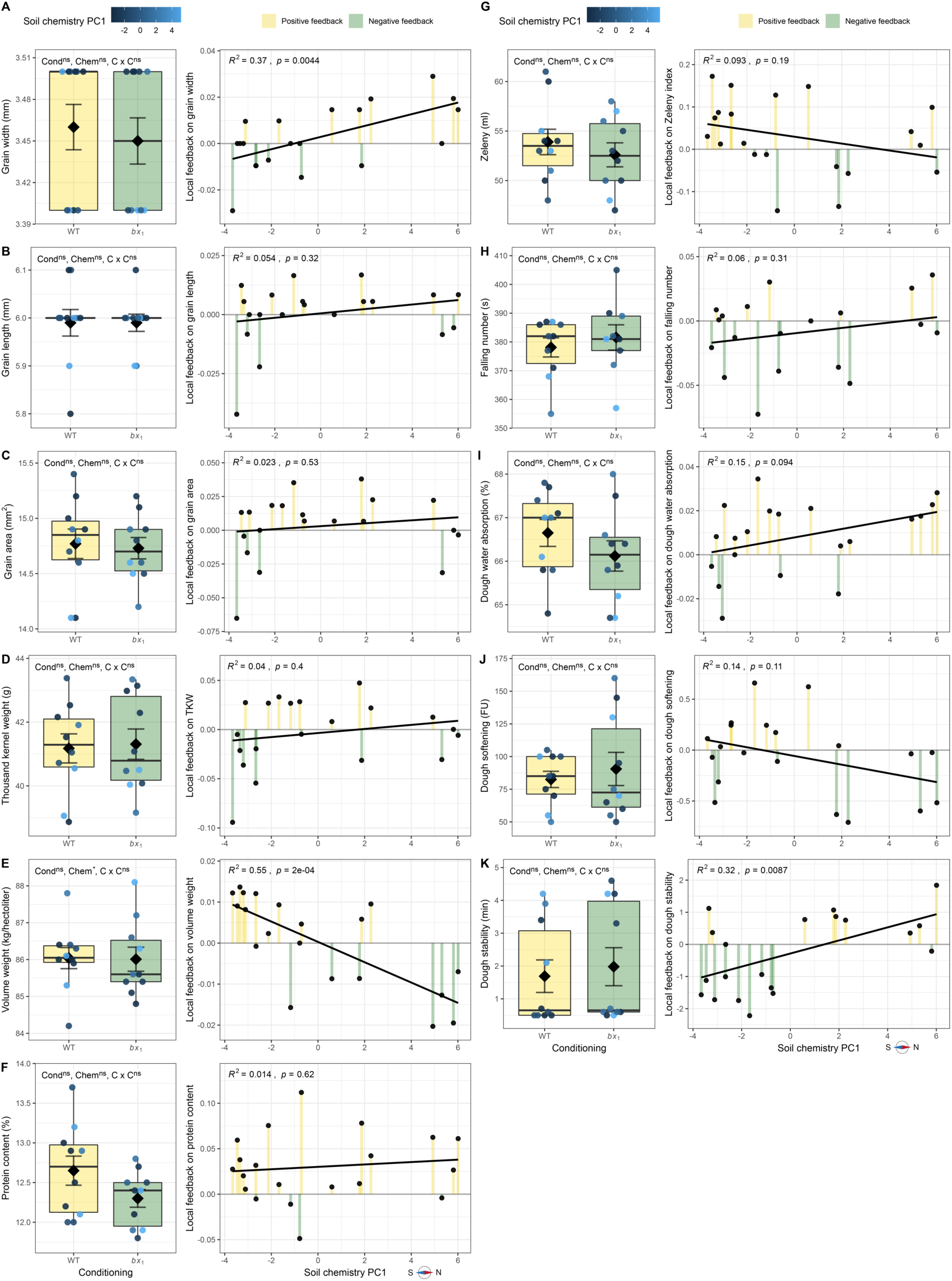
Benzoxazinoid-dependent plant-soil feedbacks on agronomic kernel parameters. For each agronomic kernel parameter boxplots (left) and local feedbacks of individual plots along the soil chemistry PC1 (right) are shown. For boxplots, phenotypes measured on plots conditioned by wild-type (WT) or benzoxazinoid-deficient *bx1* mutant maize are shown. Means ± SE and individual datapoints are included. Further, significance of ANOVA output is shown, where benzoxazinoid soil conditioning (Cond), the soil chemistry PC1 (Chem) and their interaction (C x C) were modelled. For the local feedbacks, log-response ratio values of individual plots are shown and *R^2^* and *p* value of linear regression are indicated on top. For more details on the local feedback refer to method section. Levels of significance: p < 0.001 ***, p < 0.01 **, p < 0.05 *, p > 0.05 ^ns^.

**Supplementary Figure S10.**
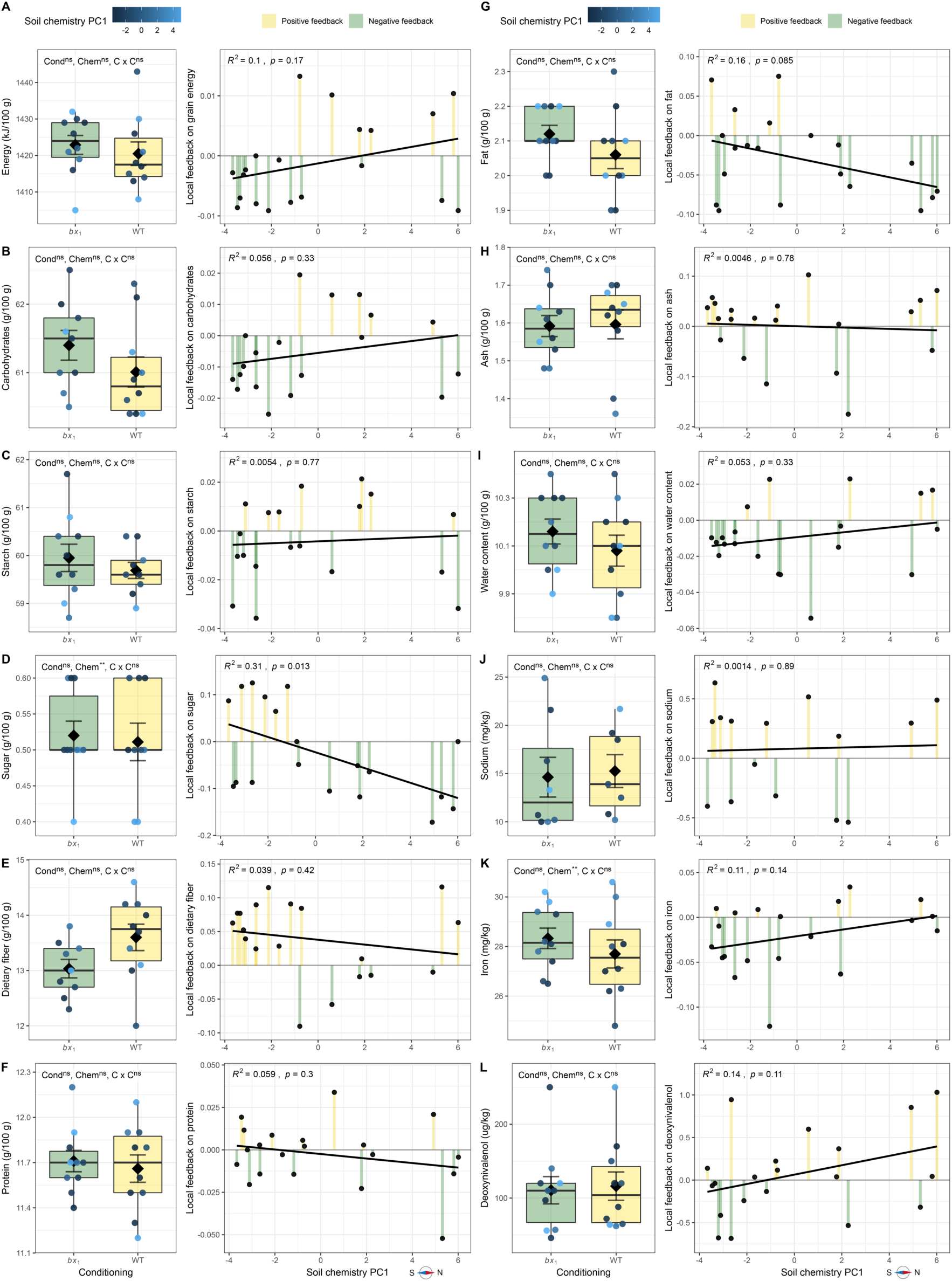
Benzoxazinoid-dependent plant-soil feedbacks on food quality parameters. Various food quality parameters were determined in the wheat kernels. For each parameter boxplots (left) and local feedbacks of individual plots along the soil chemistry (right) are shown. For boxplot, phenotypes measured on plots conditioned by wild-type (WT) or benzoxazinoid-deficient *bx1* mutant maize are shown; means ± SE and individual datapoints are included (n = 10). Further, significance of ANOVA output is shown, where benzoxazinoid soil conditioning (Cond), the soil chemistry PC1 (Chem), and their interaction (C x C) were modelled. For the local feedbacks, log-response ratio values of individual plots are shown and *R^2^* and *p* value of linear regression are indicated on top. For more details on the local feedback refer to method section. Levels of significance: p < 0.001 ***, p < 0.01 **, p < 0.05 *, p > 0.05 ^ns^.

## Supplementary Tables

**Supplementary Table S1.** **PERMANOVA of Microbiomes at maize harvest.** Compartment-wise analysis of maize genotype (WT, *bx1*), soil chemistry (PC axis 1), and their interaction for bacteria (top) and fungi(bottom). Significant *p* values are shown in bold.

| **Bacteria** |  | Df | SumOfSqs | R2 | F | Pr(>F) |
| --- | --- | --- | --- | --- | --- | --- |
| Root | Genotype | 1 | 0.2582 | 0.07514 | 2.2878 | **0.049** * |
|  | Soil chemistry PC1 | 1 | 1.2611 | 0.36695 | 11.1727 | **0.001** *** |
|  | Genotype:SoilChemPC1 | 1 | 0.1114 | 0.03241 | 0.9867 | 0.381 |
|  | Residual | 16 | 1.8060 | 0.52550 |  |  |
|  | Total | 19 | 3.4367 | 1.00000 |  |  |
| Rhizo. | Genotype | 1 | 0.22447 | 0.07741 | 1.6452 | 0.100 . |
|  | Soil chemistry PC1 | 1 | 0.82186 | 0.28342 | 6.0236 | **0.001** *** |
|  | Genotype:SoilChemPC1 | 1 | 0.21615 | 0.07454 | 1.5842 | 0.117 |
|  | Residual | 12 | 1.63728 | 0.56463 |  |  |
|  | Total | 15 | 2.89976 | 1.00000 |  |  |
| Soil | Genotype | 1 | 0.1264 | 0.03169 | 0.9343 | 0.404 |
|  | Soil chemistry PC1 | 1 | 1.6104 | 0.40374 | 11.9031 | **0.001** *** |
|  | Genotype:SoilChemPC1 | 1 | 0.0872 | 0.02186 | 0.6445 | 0.699 |
|  | Residual | 16 | 2.1646 | 0.54271 |  |  |
|  | Total | 19 | 3.9886 | 1.00000 |  |  |
| **Fungi** |  | Df | SumOfSqs | R2 | F | Pr(>F) |
| Root | Genotype | 1 | 0.32678 | 0.13110 | 3.1550 | **0.002** ** |
|  | Soil chemistry PC1 | 1 | 0.40106 | 0.16090 | 3.8721 | **0.001** *** |
|  | Genotype:SoilChemPC1 | 1 | 0.10750 | 0.04313 | 1.0379 | 0.411 |
|  | Residual | 16 | 1.65722 | 0.66487 |  |  |
|  | Total | 19 | 2.49255 | 1.00000 |  |  |
| Rhizo. | Genotype | 1 | 0.40395 | 0.15156 | 3.6085 | **0.003** ** |
|  | Soil chemistry PC1 | 1 | 0.33356 | 0.12515 | 2.9797 | **0.005** ** |
|  | Genotype:SoilChemPC1 | 1 | 0.13664 | 0.05127 | 1.2207 | 0.259 |
|  | Residual | 16 | 1.79110 | 0.67202 |  |  |
|  | Total | 19 | 2.66526 | 1.00000 |  |  |
| Soil | Genotype | 1 | 0.2436 | 0.04152 | 0.7295 | 0.944 |
|  | Soil chemistry PC1 | 1 | 0.3574 | 0.06092 | 1.0704 | 0.291 |
|  | Genotype:SoilChemPC1 | 1 | 0.2573 | 0.04386 | 0.7706 | 0.903 |
|  | Residual | 15 | 5.0089 | 0.85371 |  |  |
|  | Total | 18 | 5.8672 | 1.00000 |  |  |

Signif. codes: \*\*\* 0.001 \*\* 0.01 \* 0.05 . 0.1

**Supplementary Table S2.**
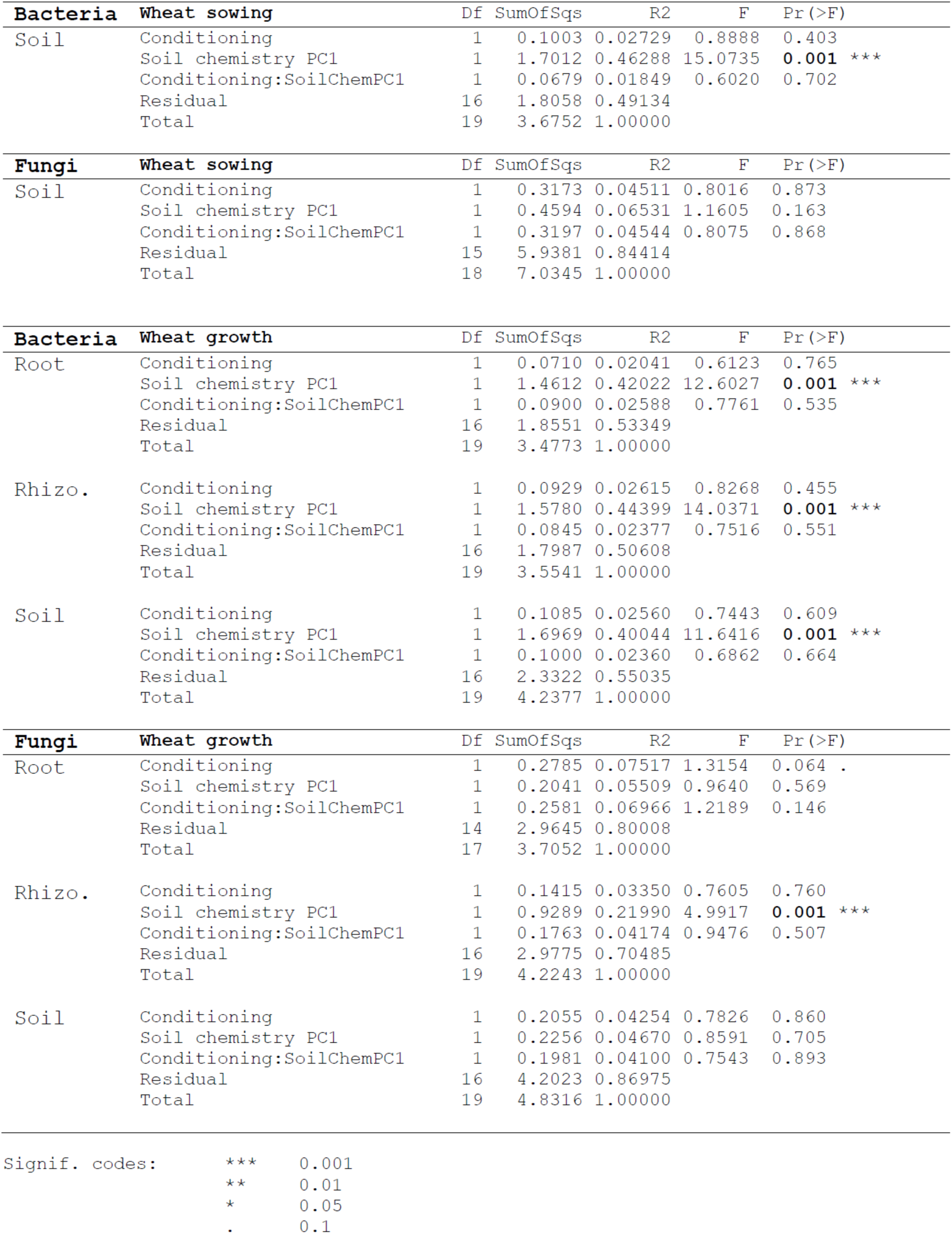
PERMANOVA of Microbiomes at wheat sowing and during wheat growth. Compartment-wise analysis of soil conditioning (WT, *bx1*), soil chemistry (PC axis 1), and their interaction for bacteria (top) and fungi(bottom) at wheat sowing and during wheat growth. Significant *p* values are shown in bold.

## Notes

### Competing Interest Statement

The authors have declared no competing interest.

## References

Andrews S. 2010.FastQC: a quality control tool for high throughput sequence data.

Benjamini Y, Hochberg Y. 1995. Controlling the False Discovery Rate: A Practical and Powerful Approach to Multiple Testing. Journal of the Royal Statistical Society: Series B (Methodological) 57: 289–300.

Bennett JA, Klironomos J. 2019. Mechanisms of plant-soil feedback: interactions among biotic and abiotic drivers. The New phytologist 222: 91–96.

Bennett JA, Maherali H, Reinhart KO, Lekberg Y, Hart MM, Klironomos J. 2017. Plant-soil feedbacks and mycorrhizal type influence temperate forest population dynamics. Science 355: 181–184.

Berendsen RL, Pieterse CMJ, Bakker PAHM. 2012. The rhizosphere microbiome and plant health. Trends in plant science 17: 478–486.

Bever JD, Platt TG, Morton ER. 2012. Microbial population and community dynamics on plant roots and their feedbacks on plant communities. Annual review of microbiology 66: 265–283.

Bodenhausen N, Horton MW, Bergelson J. 2013. Bacterial communities associated with the leaves and the roots of Arabidopsis thaliana. PLOS ONE 8: e56329.

Cadot S, Gfeller V, Hu L, Singh N, Sánchez-Vallet A, Glauser G, Croll D, Erb M, van der Heijden MGA, Schlaeppi K. 2021a. Soil composition and plant genotype determine benzoxazinoid-mediated plant-soil feedbacks in cereals. Plant, cell & environment 44: 3502–3514.

Cadot S, Guan H, Bigalke M, Walser J-C, Jander G, Erb M, van der Heijden MGA, Schlaeppi K. 2021b. Specific and conserved patterns of microbiota-structuring by maize benzoxazinoids in the field. Microbiome 9: 103.

Callahan BJ. 2018. Silva Taxonomic Training Data Formatted For Dada2 (Silva Version 132). Zenodo (DOI: 10.5281/zenodo.1172783).

Callahan BJ, McMurdie PJ, Rosen MJ, Han AW, Johnson AJA, Holmes SP. 2016. DADA2: High-resolution sample inference from Illumina amplicon data. Nature methods 13: 581–583.

Chelius MK, Triplett EW. 2001. The Diversity of Archaea and Bacteria in Association with the Roots of Zea mays L. Microbial ecology 41: 252–263.

Choma M, Tahovská K, Kaštovská E, Bárta J, Růžek M, Oulehle F. 2020. Bacteria but not fungi respond to soil acidification rapidly and consistently in both a spruce and beech forest. FEMS microbiology ecology 96: fiaa174.

Cotton TEA, Pétriacq P, Cameron DD, Meselmani MA, Schwarzenbacher R, Rolfe SA, Ton J. 2019. Metabolic regulation of the maize rhizobiome by benzoxazinoids. The ISME Journal 13: 1647–1658.

Etzerodt T, Mortensen AG, Fomsgaard IS. 2008. Transformation kinetics of 6-methoxybenzoxazolin-2-one in soil. Journal of environmental science and health. Part. B, Pesticides, food contaminants, and agricultural wastes 43: 1–7.

Forero LE, Grenzer J, Heinze J, Schittko C, Kulmatiski A. 2019. Greenhouse- and Field-Measured Plant-Soil Feedbacks Are Not Correlated. Frontiers in Environmental Science 7: 184.

Fry EL, Johnson GN, Hall AL, Pritchard WJ, Bullock JM, Bardgett RD. 2018. Drought neutralises plant-soil feedback of two mesic grassland forbs. Oecologia 186: 1113–1125.

Gardes M, Bruns TD. 1993. ITS primers with enhanced specificity for basidiomycetes--application to the identification of mycorrhizae and rusts. Molecular ecology 2: 113–118.

Gfeller V, Thönen L, Erb M. 2023. Root-exuded secondary metabolites can alleviate negative plant-soil feedbacks. bioRxiv. doi: 10.1101/2023.04.09.536155.

Gfeller V, Waelchli J, Pfister S, Deslandes-Hérold G, Mascher F, Glauser G, Aeby Y, Mestrot A, Robert CA, Schlaeppi K et al. 2022. Plant secondary metabolite-dependent plant-soil feedbacks can improve crop yield in the field. bioRxiv. doi: 10.1101/2022.11.09.515047.

Hannula SE, Heinen R, Huberty M, Steinauer K, Long JR de, Jongen R, Bezemer TM. 2021. Persistence of plant-mediated microbial soil legacy effects in soil and inside roots. Nature Communications 12: 5686.

Hu L, Mateo P, Ye M, Zhang X, Berset JD, Handrick V, Radisch D, Grabe V, Köllner TG, Gershenzon J et al. 2018a. Plant iron acquisition strategy exploited by an insect herbivore. Science 361: 694–697.

Hu L, Robert CAM, Cadot S, Zhang X, Ye M, Li B, Manzo D, Chervet N, Steinger T, van der Heijden MGA et al. 2018b. Root exudate metabolites drive plant-soil feedbacks on growth and defense by shaping the rhizosphere microbiota. Nature communications 9: 2738.

Hu L, Wu Z, Robert CAM, Ouyang X, Züst T, Mestrot A, Xu J, Erb M. 2021. Soil chemistry determines whether defensive plant secondary metabolites promote or suppress herbivore growth. Proceedings of the National Academy of Sciences 118: e2109602118.

Huang AC, Jiang T, Liu Y-X, Bai Y-C, Reed J, Qu B, Goossens A, Nützmann H-W, Bai Y, Osbourn A. 2019. A specialized metabolic network selectively modulates Arabidopsis root microbiota. Science 364: eaau6389.

Kaisermann A, Vries FT de, Griffiths RI, Bardgett RD. 2017. Legacy effects of drought on plant-soil feedbacks and plant-plant interactions. The New phytologist 215: 1413–1424.

Kos M, Tuijl MAB, Roo J de, Mulder PPJ, Bezemer TM. 2015a. Plant-soil feedback effects on plant quality and performance of an aboveground herbivore interact with fertilisation. Oikos 124: 658–667.

Kos M, Tuijl MAB, Roo J de, Mulder PPJ, Bezemer TM. 2015b. Species-specific plant-soil feedback effects on above-ground plant-insect interactions. Journal of Ecology 103: 904–914.

Kudjordjie EN, Sapkota R, Steffensen SK, Fomsgaard IS, Nicolaisen M. 2019. Maize synthesized benzoxazinoids affect the host associated microbiome. Microbiome 7: 59.

Kuerban M, Cong W-F, Jing J, Bezemer TM. 2022. Microbial soil legacies of crops under different water and nitrogen levels determine succeeding crop performance. Plant and Soil. doi: 10.1007/s11104-022-05412-6.

Lê S, Josse J, Husson F. 2008. FactoMineR : An R Package for Multivariate Analysis. Journal of Statistical Software 25: 1–18.

Long JR de, Fry EL, Veen GF, Kardol P. 2019. Why are plant–soil feedbacks so unpredictable, and what to do about it? Functional Ecology 33: 118–128.

Ma H-K, Pineda A, van der Wurff AWG, Raaijmakers C, Bezemer TM. 2017. Plant-Soil Feedback Effects on Growth, Defense and Susceptibility to a Soil-Borne Disease in a Cut Flower Crop: Species and Functional Group Effects. Frontiers in plant science 8: 2127.

Macías FA, Oliveros-Bastidas A, Marín D, Castellano D, Simonet AM, Molinillo JMG. 2004. Degradation studies on benzoxazinoids. Soil degradation dynamics of 2,4-dihydroxy-7-methoxy-(2H)-1,4-benzoxazin-3(4H)-one (DIMBOA) and its degradation products, phytotoxic allelochemicals from gramineae. Journal of agricultural and food chemistry 52: 6402–6413.

Maresh J, Zhang J, Lynn DG. 2006. The innate immunity of maize and the dynamic chemical strategies regulating two-component signal transduction in Agrobacterium tumefaciens. ACS chemical biology 1: 165–175.

Mariotte P, Mehrabi Z, Bezemer TM, Deyn GB de, Kulmatiski A, Drigo B, Veen GFC, van der Heijden MGA, Kardol P. 2018. Plant-Soil Feedback: Bridging Natural and Agricultural Sciences. Trends in ecology & evolution 33: 129–142.

Martin M. 2011. Cutadapt removes adapter sequences from high-throughput sequencing reads. EMBnet.journal 17: 10.

McMurdie PJ, Holmes S. 2013. phyloseq: an R package for reproducible interactive analysis and graphics of microbiome census data. PLOS ONE 8: e61217.

Nannipieri P, Kandeler E, Ruggiero P. 2002. Enzyme activities and microbiological and biochemical processes in soil. in Enzymes in the environment. New York, NY: Marcel Dekker.

Niemeyer HM. 2009. Hydroxamic acids derived from 2-hydroxy-2H-1,4-benzoxazin-3(4H)-one: key defense chemicals of cereals. Journal of agricultural and food chemistry 57: 1677–1696.

Nilsson RH, Larsson K-H, Taylor AFS, Bengtsson-Palme J, Jeppesen TS, Schigel D, Kennedy P, Picard K, Glöckner FO, Tedersoo L et al. 2019. The UNITE database for molecular identification of fungi: handling dark taxa and parallel taxonomic classifications. Nucleic acids research 47: D259–D264.

Oksanen J, Blanchet FG, Friendly M, Kindt R, Legendre P, McGlinn D, Minchin PR, O’Hara RB, Simpson GL, Solymos P et al. 2020. vegan: Community Ecology Package. [WWW document] URL https://CRAN.R-project.org/package=vegan.

Pang Z, Chen J, Wang T, Gao C, Li Z, Guo L, Xu J, Cheng Y. 2021. Linking Plant Secondary Metabolites and Plant Microbiomes: A Review. Frontiers in plant science 12: 621276.

Pieterse CMJ, Zamioudis C, Berendsen RL, Weller DM, van Wees SCM, Bakker PAHM. 2014. Induced systemic resistance by beneficial microbes. Annual review of phytopathology 52: 347–375.

Pineda A, Kaplan I, Hannula SE, Ghanem W, Bezemer TM. 2020. Conditioning the soil microbiome through plant–soil feedbacks suppresses an aboveground insect pest. The New phytologist 226: 595–608.

Quast C, Pruesse E, Yilmaz P, Gerken J, Schweer T, Yarza P, Peplies J, Glöckner FO. 2013. The SILVA ribosomal RNA gene database project: improved data processing and web-based tools. Nucleic acids research 41: D590–D596.

R Core Team. 2021. R: A Language and Environment for Statistical Computing. Vienna, Austria. [WWW document] URL https://www.R-project.org/.

Revillini D, Gehring CA, Johnson NC. 2016. The role of locally adapted mycorrhizas and rhizobacteria in plant–soil feedback systems. Functional Ecology 30: 1086–1098.

Rousk J, Bååth E, Brookes PC, Lauber CL, Lozupone C, Caporaso JG, Knight R, Fierer N. 2010. Soil bacterial and fungal communities across a pH gradient in an arable soil. The ISME Journal 4: 1340–1351.

Schandry N, Becker C. 2020. Allelopathic Plants: Models for Studying Plant-Interkingdom Interactions. Trends in plant science 25: 176–185.

Schittko C, Runge C, Strupp M, Wolff S, Wurst S. 2016. No evidence that plant-soil feedback effects of native and invasive plant species under glasshouse conditions are reflected in the field. Journal of Ecology 104: 1243–1249.

Smith-Ramesh LM, Reynolds HL. 2017. The next frontier of plant–soil feedback research: unraveling context dependence across biotic and abiotic gradients. Journal of Vegetation Science 28: 484–494.

Strebel S, Levy Häner L, Mattin M, Schaad N, Morisoli R, Watroba M, Girard M, Courvoisier N, Berberat J, Grandgirard R et al. 2022. Liste der empfohlenen Getreidesorten für die Ernte 2023. Agroscope Transfer 443: 1–8.

Stringlis IA, Jonge R de, Pieterse CMJ. 2019. The Age of Coumarins in Plant-Microbe Interactions. Plant & cell physiology 60: 1405–1419.

Tawaha AM, Turk MA. 2003. Allelopathic Effects of Black Mustard (Brassica nigra) on Germination and Growth of Wild Barley (Hordeum spontaneum). Journal of Agronomy and Crop Science 189: 298–303.

Tzin V, Fernandez-Pozo N, Richter A, Schmelz EA, Schoettner M, Schäfer M, Ahern KR, Meihls LN, Kaur H, Huffaker A et al. 2015. Dynamic Maize Responses to Aphid Feeding Are Revealed by a Time Series of Transcriptomic and Metabolomic Assays. Plant physiology 169: 1727–1743.

van der Putten WH, Bardgett RD, Bever JD, Bezemer TM, Casper BB, Fukami T, Kardol P, Klironomos JN, Kulmatiski A, Schweitzer JA et al. 2013. Plant-soil feedbacks: the past, the present and future challenges. Journal of Ecology 101: 265–276.

Voges MJEEE, Bai Y, Schulze-Lefert P, Sattely ES. 2019. Plant-derived coumarins shape the composition of an Arabidopsis synthetic root microbiome. Proceedings of the National Academy of Sciences of the United States of America 116: 12558–12565.

Weiss S, Xu ZZ, Peddada S, Amir A, Bittinger K, Gonzalez A, Lozupone C, Zaneveld JR, Vázquez-Baeza Y, Birmingham A et al. 2017. Normalization and microbial differential abundance strategies depend upon data characteristics. Microbiome 5: 27.

White TJ, Bruns T, Lee S, Taylor J. 1990. AMPLIFICATION AND DIRECT SEQUENCING OF FUNGAL RIBOSOMAL RNA GENES FOR PHYLOGENETICS. In: PCR Protocols. Elsevier, 315– 322.

Wickham H, Averick M, Bryan J, Chang W, McGowan L, François R, Grolemund G, Hayes A, Henry L, Hester J et al. 2019. Welcome to the Tidyverse. Journal of Open Source Software 4: 1686.

Yu P, He X, Baer M, Beirinckx S, Tian T, Moya YAT, Zhang X, Deichmann M, Frey FP, Bresgen V et al. 2021. Plant flavones enrich rhizosphere Oxalobacteraceae to improve maize performance under nitrogen deprivation. Nature plants 7: 481–499.

Zhao Z, Gao X, Ke Y, Chang M, Xie L, Li X, Gu M, Liu J, Tang X. 2019. A unique aluminum resistance mechanism conferred by aluminum and salicylic-acid-activated root efflux of benzoxazinoids in maize. Plant and Soil 437: 273–289.

Zhou S, Richter A, Jander G. 2018. Beyond Defense: Multiple Functions of Benzoxazinoids in Maize Metabolism. Plant & cell physiology 59: 1528–1537.

